# Prediction of Kv11.1 potassium channel PAS-domain variants trafficking via machine learning

**DOI:** 10.1101/2021.11.03.467212

**Authors:** Kalyan Immadisetty, Xuan Fang, Cassandra M. Hartle, Thomas P. McCoy, Regeneron Genetics Center, Tooraj Mirshahi, Brian P. Delisle, Peter M. Kekenes-Huskey

## Abstract

Congenital long QT syndrome (LQTS) is characterized by a prolonged QT-interval on an electrocardiogram (ECG). An abnormal prolongation in the QT-interval increases the risk for fatal arrhythmias despite otherwise normal metrics of cardiac function. Genetic variants in several different cardiac ion channel genes, including *KCNH2*, are known to cause LQTS. The population frequency of rare non-synonymous (missense) variants in LQTS-linked genes far outpaces the true incidence of the disease. Therefore, only a small percentage of missense variants identified in LQTS-linked genes are expected to associate with LQTS. Because of a lack of clear association between variants identified in LQTS-linked alleles and diseases, most variants are classified as variants of uncertain physiological significance (VUS). Here, we evaluated whether structure-based molecular dynamics (MD) simulations and machine learning (ML) can improve the identification of missense variants in LQTS-linked genes that associate with LQTS. To do this, we focused on investigating *KCNH2* missense variants in the Kv11.1 channel protein shown to have wild type (WT) like or loss-of-function (LOF) phenotypes *in vitro*. We focused on *KCNH2* missense variants that disrupt normal Kv11.1 channel protein trafficking, as it is the most common LOF phenotype for LQTS-associated variants. Specifically, we used these computational techniques to correlate structural and dynamic changes in the Kv11.1 channel protein PAS domain (PASD) with Kv11.1 channel protein trafficking phenotypes. These simulations unveiled several molecular features, including the numbers of hydrating waters and H-Bonds, as well as FoldX scores, that are predictive of trafficking. We then used statistical and ML (Decision tree (DT), Random forest (RF), and Support vector machine (SVM)) techniques to classify variants using these simulation-derived features. Together with bioinformatics data, such as sequence conservation and folding energies, we were able to predict with reasonable accuracy (≈75%) which *KCNH2* variants do not traffic normally. We conclude, structure-based simulations of *KCNH2* variants localized to the Kv11.1 channel PASD led to a significant improvement (≈10%) in classification accuracy and this approach should therefore be considered to complement the classification of VUS in the Kv11.1 channel PASD.

## 2 Introduction

Every year congenital Long QT syndrome (LQTS) kills thousands. This is not because LQTS is difficult to treat, but it is difficult to diagnose early before a deadly arrhythmia occurs. Ideally genetic screening could facilitate the early identification and prophylactic treatment of LQTS. This “gene-first” approach has the potential to become a reality in the era of Precision Medicine.[Shah2016; NRC2011] However, genetic testing often identifies novel, rare missense variants of uncertain physiological significance (VUS).[kapa2009; Taggart2007; prio2013; Schwartz2013; Crotti2008]

The number of VUS identified in LQTS genes grows as more people are genotyped. Federally funded Precision Medicine initiatives, including NIH’s All of Us Research Program and the Department of Veterans Affairs Million Veteran Program, are building comprehensive genetic biobanks for millions of participants. Hospitals are now starting to generate their own genetic biobanks. For example, Geisinger Health’s MyCode® initiative has already developed a system that directly links the Whole Exome Sequence (WES) data for over 100,000 patients to their electronic health records (EHR). Moreover, the decreasing cost of genetic testing is stimulating FDA-approved direct-to-consumer genetic healthcare reports.[Check2017] The challenge we now face is how to reliably determine which VUS in LQTS genes are dysfunctional and potentially disease causing.[kapa2009]

In order to prevent the misdiagnosis of patients based primarily on genetic test for a VUS, the current clinical recommendations now limit genetic screening to phenotypically positive patients.[Ackerman2011; prio2013] This “phenotype-first” approach prevents the early identification of LQTS patients starting from the increasingly available patient genetic data (e.g. WES).[kapa2009] Since one of the presenting symptoms of LQTS is sudden death, it is important to develop reliable gene-first approaches that identify at risk patients before LQTS strikes. The goal of this study was to identify and develop strategies that reliably clarify the physiological significance of LQTS-linked VUS. In this study we focus on LQTS type 2 (LQT2), which is caused by loss-of-function (LOF) mutations in the *KCNH2*-encoding voltage-gated K^+^ channel *α*-subunit (Kv11.1). Previous studies show most LQT2-linked missense mutations disrupt the normal intracellular transport (trafficking) of the Kv11.1 channel protein to the cell surface membrane [Anderson2014; Anderson2020]. Many of these trafficking deficient LQT2-linked mutations localize to the N-terminal Per-Arnt-Sim domain (PASD). Trafficking-deficient LQT2-linked mutations in the Kv11.1 channel PASD decrease PASD solubility. These findings suggest that LOF LQT2-causing variants perturb the normal channel domain structure. Since structural data are available for the PAS domain and structure-based molecular simulations are commonly used for predicting impacts of missense mutations on protein function, this raises the intriguing possibility that computer simulation analyses can be used to reliably predict which Kv11.1 VUS negatively affect Kv11.1 channel trafficking.

We tested the hypothesis that sequence-and structure-based computational analyses could reliably predict whether trafficking-deficient, LOF PASD variants tended to perturb the native folding of the PASD structure. To investigate this hypothesis, we performed bioinformatics analyses of the PASD primary sequence in complement with structure-based analyses and simulations.

The structural approach comprised folding free energy scoring via FOLDX, analyses of domain solubility via APBS, and molecular dynamics (MD) simulations of the Kv11.1 channel PASD structure that was solved at 1.96 Å resolution by Tang *et al*. using X-ray crystallography [tang2016]. With this approach, we considered the WT and 74 PASD-localized Kv11.1 channel protein variants whose solubility was assayed by Anderson *et al*. [Anderson2020]. From these simulations, we assessed the PASD’s native structure, conformational dynamics, and several physiochemical properties (e.g., hydrophobicity) of the PASD. Changes in these PASD properties were evaluated to determine their capacity to discriminate trafficking from non-trafficking Kv11.1 channel protein variants.

These MD-derived properties were used with statistical and machine learning (ML) techniques to classify variants as soluble or insoluble, and this was used as a proxy for normal and LOF trafficking phenotypes. Despite the strong bias of the training data toward variants with impaired trafficking, we found that the ML approaches, and in particular the random forest, achieved higher accuracy using bioinformatics-based features alone. Using the trained ML models, we proposed and assessed a list of novel variants (S26G, R27L, N33T, R35L, C39R, V41I, D46E, D46N, C49S, R76L, R76H, A79T, L87R, R92H, R92C, F98L, L107P, V115M, N117S, D119H, V131A, V131L, M133T) that we predicted would yield normal and LOF phenotypes. Nearly all variants were predicted to traffic. Electrocardiograph data collected from patients harboring these mutations did not present obvious prolongation of the QT interval, which is consistent with our predictions of normal trafficking. Altogether, this study advocates for including structure-based simulation methods and machine learning strategies to facilitate the assignment of *KCNH2* VUS.

## 3 Results

### 3.1 Introduction to characterized variants

- Roadmap of known variants
- Loose description of general characteristics (wt-like, correctable, etc)
- Classification by ‘conceptual’ phenomena Our foldx and comparison against other solubility data

### 3.2 Predictive features of PASD solubility

#### 3.2.1 Bioinformatics analyses of the entire variant set

We demonstrated in Section 3.1 that PASD variants thus far do not exhibit obvious patterns linking trafficking to intuitive features like proximity to known features, charge etc. However, leveraging the approach from [Anderson2020] *et al*., we confirm that the solubility, a necessary conditionfor trafficking, correlates with the folding free energy of a variant relative to WT. Given the ease of computing free energies via FOLDX, we next evaluated a series of other easy-to-compute metrics that can be gleaned from either the protein sequence or its 3D structure. Scores obtained from theses evaluates are compared to the relative solublity measured by Anderson *et al*. [Anderson2020]

We first considered ‘bioinformatics-based’ criteria such as the amino acid side chain hydrophobicity and volume, conservation scores for each sequence position, and FoldX scores for a given missense mutation (45 trafficking and 30 non-trafficking variants in total). The variants’ raw values for each criterion were offset by the corresponding value for the WT Kv11.1 channel PASD, such that the wild-type sequence yielded a score of zero. The first two criteria were computed using the Kv11.1 channel protein primary sequence. The hydrophobicity changes generally assign more negative values for apolar to charged amino acids, less negative values for variants from apolar to polar amino acids, and positive values for charged to charged to apolar substitutions. Similarly, we include amino acid volume changes to reflect variants from large (e.g., W, Y, and F) side groups into smaller groups (e.g. A). These features are intended to identify variants that we postulated could disrupt the normal packing of the PASD globular structure and promote unfolding. The conservation score reflects the degree to which a given amino acid site is evolutionarily conserved; conservation scores 1 and 9 are considered highly variable and conserved, respectively. The Foldx scores, meanwhile, utilize an all-atom force field-based empirical scoring function [SCHY2005] to estimate the impact of a given variant on the free energy, *G*, of the mutated protein relative to the WT.

Hence, Δ*G >* 0 signifies a destabilizing mutation. We rank order the variants according to these features in Fig. 3. The WT-shifted raw feature values are presented on the left axes, while the right axes report WT-shifted z-scores as described in Section 6.3.1 The scores corresponding to the WT are 0 in each panel. All variants are highlighted as normally trafficking versus LOF variants with green and red bars, respectively. In the event that each classifier were perfect, LOF variants would have scores that are distinct (e.g. positive) from those of the WT and with normal trafficking (e.g. *z*_*j*_ ≤ 0). As an example, the FoldX scores for non-traffickers tend to have z-scores above 0.5, with a few traffickers adopting similar values. In general, the Foldx scores tend to better partition the trafficking from non-trafficking, with the latter yielding more positive z-scores; however, the features was imperfect.

**Figure 1:**
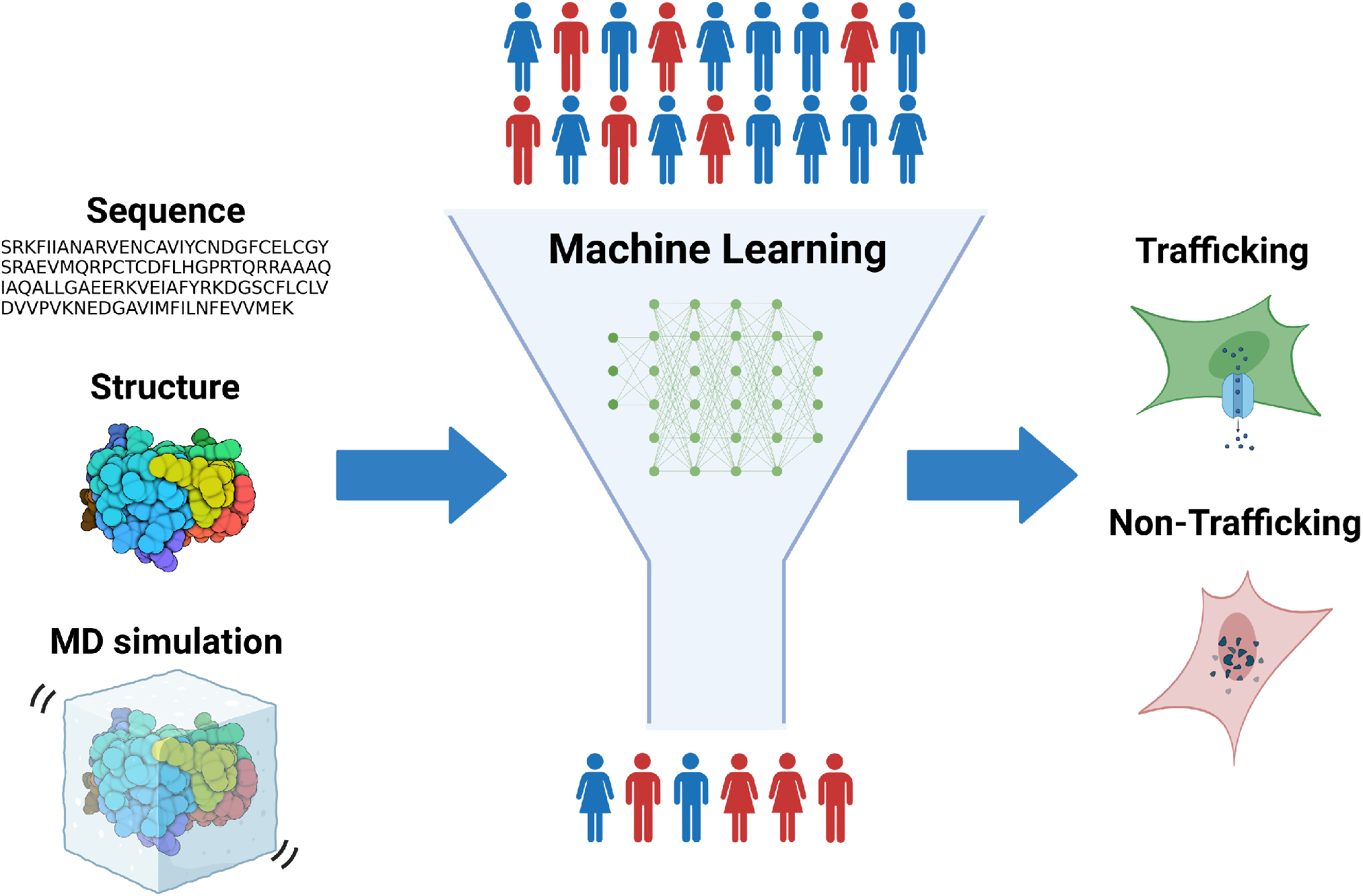
Schematic of study. We developed a strategy for identifying potential Kv11.1 loss-of-function phenotypes for patients that present missense mutations within the Per-Arnt-Sim domain sequence of KCNH2. Our strategy utilizes several machine learning methods to classify variants into class II (LOF) and non-class II phenotypes using features from the primary sequence, three-dimensional structures, and molecular dynamics (MD) simulations. https://app.biorender.com/illustrations/62757a31cafe185efeaf6d47

**Figure 2:**
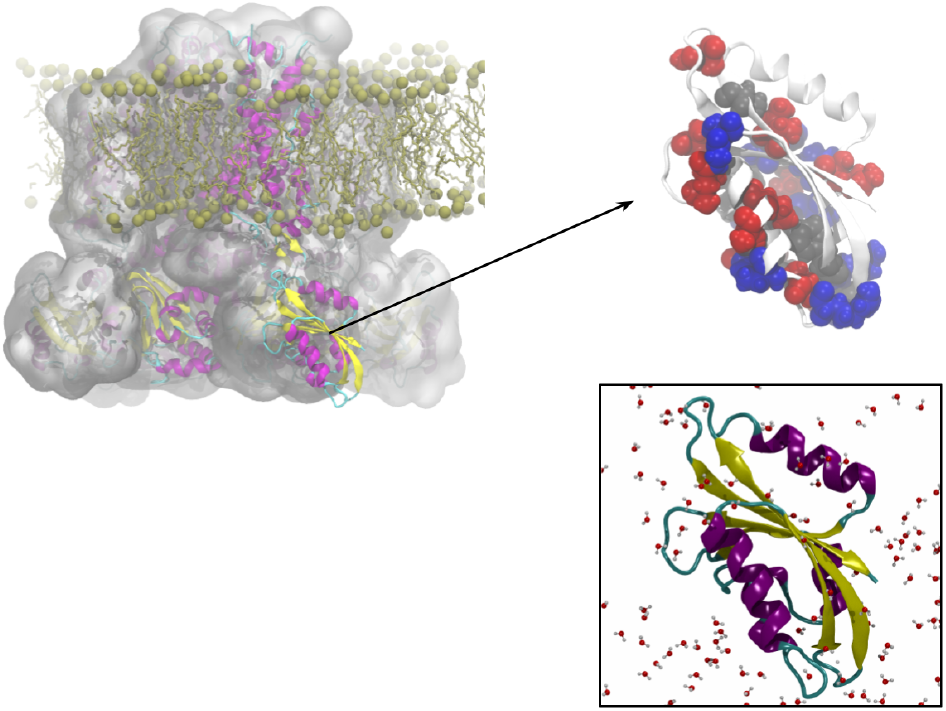
Schematic of the Kv11.1 potassium channel study. The Kv11.1 protein comprises mainly of three domains: transmembrane domain, PAS domain and cyclic nucleotide binding domain (see upper left). Structure of PASD from PDB: 4HQA [tang2016] with class II variants marked in red, non-class II as blue, and gray represents variants with both behaviors (upper right). Bottom panel depicts the isolated and solvated PASD used for molecular dynamics simulations.

**Figure 3:**
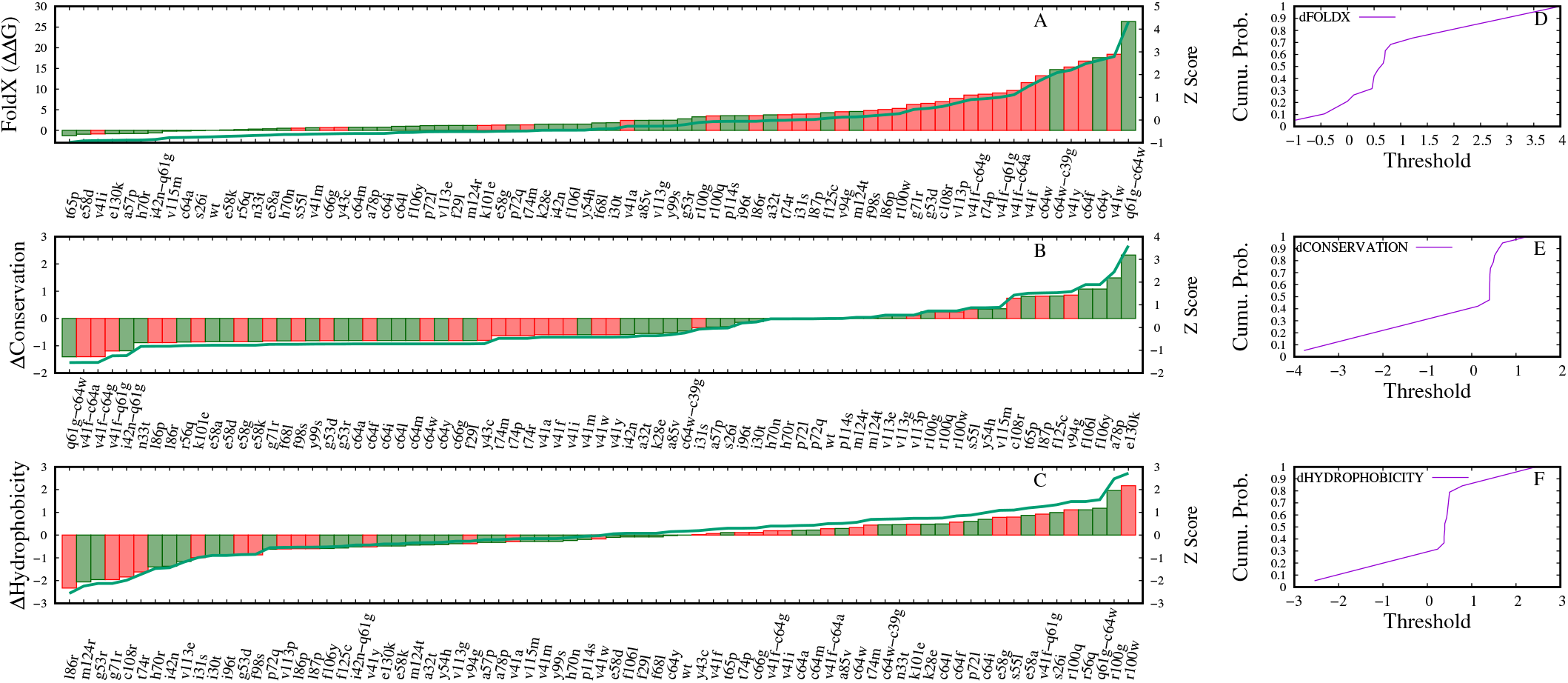
Ranking of variants by bioinformatics scores. Shown in the left panels are the changes of respective bioinformatics scores of the variants with respect to the WT. Y2-axes reflect the Z-score, which describes the distribution of the bioinformatics score of a given variant compared to the population mean. Green and red represent trafficking and non-trafficking variants, respectively. Right panels represent cumulative probabilities as a function of threshold values.

To quantitatively assess the ability to differentiate normal (non-class II) and LOF (class II), we compute the conditional probability for LOF variants to adopt a z-score above some threshold value *λ*. These data are summarized in the second column of Fig. S7 as the class II (CII) conditional probability, *p*_*ij*_(*CII* | *z*_*ij*_ *> λ*_*j*_) (see Eq. 1 in Section 6.3.1) that we abbreviate as *p*_*j*_ for convenience. With these data, we observe that *p*_*j*_ for the FoldX mostly monotonically approaches an 80% classification rate until *z*_*i*_ ≈ 0, after which the probability approaches 0. In other words, variants with Foldx z-scores near 0 have an 80% true positive rate of classification as a LOF variant. Meanwhile, the *p*_*j*_ values based on the conservation scores also monotonically increase, but are maximized at 65%. In contrast, less predictive metrics such as the hydrophobicity score yielded *p*_*j*_ curves that were nonmonotonic and roughly 50% in scale. Interestingly, the amino acid volume score also exhibited a nonmonotonic trend. This contrasts with what has been reported in several studies, where amino acid volume is correlated with LOF. The features we considered are generally uncorrelated (see Fig. S1). Hence, the P(LOF) suggest Foldx has the most predictive power for differentiating the non-traffickers from traffickers. Later we will use these probabilities to classify variants.

#### 3.2.2 Molecular dynamics simulations of variants

We next performed MD simulations on the 75 PASD variants. This was done to determine if the missense variants impacted the structure, dynamics, and energetics of the Kv11.1 channel PASD to yield additional features that could be predictive of LOF phenotypes. The features we considered include solvent accessible surface area (SASA), number of hydrating waters, number of hydrogen bonds, helicity, beta character, turns, coils, 3-10 helix, root mean squared deviations (RMSD), root mean squared fluctuations (RMSF) and solvation energy computed by APBS. The SASA, hydrating waters and solvation energy criteria are intended to assess changes in protein solubility. The number of hydrogen bonds and secondary structure metrics (helicity, 3-10 helix, etc.), reflect changes in protein folding. RMSD and RMSF assess changes in a proteins dynamic motions. In Fig. S2 we present rank ordered data for each variant with the most predictive features.

As an improvement over the bioinformatics-based features, the structure-based features yielded *p*_*j*_ → 100% for almost all features we considered. We observed that *p*_*j*_ for the hydrating water feature was mostly monotonic, e.g. LOF variants tend to have more hydrating waters than WT reference. We additionally observed that LOF variants tend to have a lower number of hydrogen bonds compared to WT (− Δ*Waters*) and are correlated (see Fig. S2) with nearly all variants with normal trafficking restricted to the low water count, high hydrogen bond values. We anticipate that the increased number of waters in the LOF variants is attributed to the partial unfolding of the PAS structure. This apparent unfolding coincides with the variant structures having fewer hydrogen bonds than the WT. Hence, these features may improve classification beyond bioinformatics-based scores alone. Other metrics exhibited significant nonmonotonic behavior, such as for the *SASA* and *coils* features. This nonmonotonic behavior is expected to complicate the classification of variants.

To provide context for the MD features we assessed in the previous section, we present representative structural and dynamic data for the WT, the T74P LOF variant, and a normally trafficking, V113E. In Fig. S5 we superimpose three representative structures obtained from these simulations. Each variant presents a similar backbone conformation relative to the WT, which suggests all adopt roughly the same global fold. We next evaluate the features identified in the previous section as a potential basis for LOF versus normal phenotypes. We first examined changes in the hydrogen binding networks intrinsic to the WT structure relative to the site-directed variants. The WT variant features 66 hydrogen bonds, relative to about 65 for V113E and 62 for T74P. In Fig. S6 we highlight where the loss of hydrogen bonding occurs for the T74P variant. As anticipated, the hydrogen bond loss occurs near the site of mutation. The hydrogen bond interactions are found between T74 and other residues such as H70, G71, and I96 in the WT (Fig. S6A) are lost in the T74P variant (Fig. S6B)

We also observe that the root mean square fluctuations are generally low for the WT and V113E, with notable increases at residues 35–40, 87–93, and 117–123, which comprise loops spanning folded beta sheets and helices. In contrast, for the T74P variant, we observe both a substantial increase in RMSF localized to the site 74 region (Fig. S5) as well as a general increase in the baseline RMSF across all residues. We were unable to directly relate these structural and dynamic changes to differences in each variant’s hydrating water shell. As is made clear in the conditional probabilities plots shown in Fig. S2, the variants have diverse effects on the MD-derived features that can have both local and global impacts on protein structure and dynamics.

### 3.3 Classifiers

In the previous section, we identified sequence-and structure-based features that are moderately predictive of normal-versus LOF phenotypes. However, the significant overlap for the non-trafficking and trafficking in these features limits a straightforward classification scheme. In Section 3.3, we leverage these data in aggregate to predict a variant’s phenotype. We approach this aim by training and evaluating several classifiers that use these features to predict the phenotype in a test data set. Per standard practice guidelines in [xu2018], we partition our data into a training set (70%, 52 entries) and a test set (30%, 23 entries). These classifiers include a probabilistic classifier, decision tree, random forests, and support vector machine, which are generally well-suited for small data sets [decision2020]. Performance metrics defined in Section 6.4 are included in these evaluations.

#### 3.3.1 Probabilistic classifier

We first construct a classifier based on the product of the conditional probabilities estimated above. This classifier assumes that each feature is statistically independent, although we recognize that some features are correlated (see Fig. S3). Variants for which *P*_*i*_(*CII* | {*z*_*i*1_, … *z*_*in*_}), the left-hand side of Eq. 2, exceeds a user-defined Λ are classified as non-trafficking. This probability is annotated as *P*_*i*_ for clarity. With this approach, we present a receiver operator characteristic (ROC) curve in Fig. S7. These ROC curves enumerate the true positives, e.g., the number of CII variants versus the total number of variants with *P*_*i*_ *>* Λ, as well as the false positive rate, e.g., the number of CII variants with *P*_*i*_ ≤ Λ. Random features that are unable to discriminate true and false positives, i.e., yield equal rates for both terms as a function of the threshold, are designated by the dashed diagonal black line. Metrics that perform better than random exhibit ROC curves above this line, which is quantified by reporting the area under the curve (AUC) values. AUC values approaching 1.0 represent metrics that maximize the TPR and minimize the FPR. Using the ROC curves, we evaluated the performance of the classifier on three different feature sets: MD alone, bioinformatics alone, and MD+bioinformatics (all) features. The results indicate that with the MD and all feature sets, the classifier achieves good true positive rates relative to false positive rates, as evidenced by an AUC of 0.75. As an example, many of the threshold values (Γ) in Fig. S7A achieve TPRs within 0.6 and 1.0 and have FPRs of less than 0.4. This indicates that the probabilities are reasonable classifiers for partitioning the variants into CII or non-class II ((-)CII) classifications based on threshold alone. We additionally provide a summary of performance metrics in Table 1 based on statistics collected from 500 bootstrapped samples. These metrics include accuracy, precision, recall, specificity, and F1-score. Of these, we focus on the precision, recall, and F1-score metrics in our analyses. The precision reflects the probability of correctly classifying positive hits, while recall assesses the completeness of the classification, e.g., classifying all positives in the data set. Our analyses indicate that the probabilistic classifier yields precisions of 65 for both CII and (-)CII while the recall is 24 and 93. The F1 score assesses the trade-off in recall and precision performance; our strategy therefore yielded F1-scores of 33 and 76 for the CII and (-)CII classifications, respectively. Overall, the probabilistic classifier performed well with classifying variants as class II and non-class II

**Table 1:**
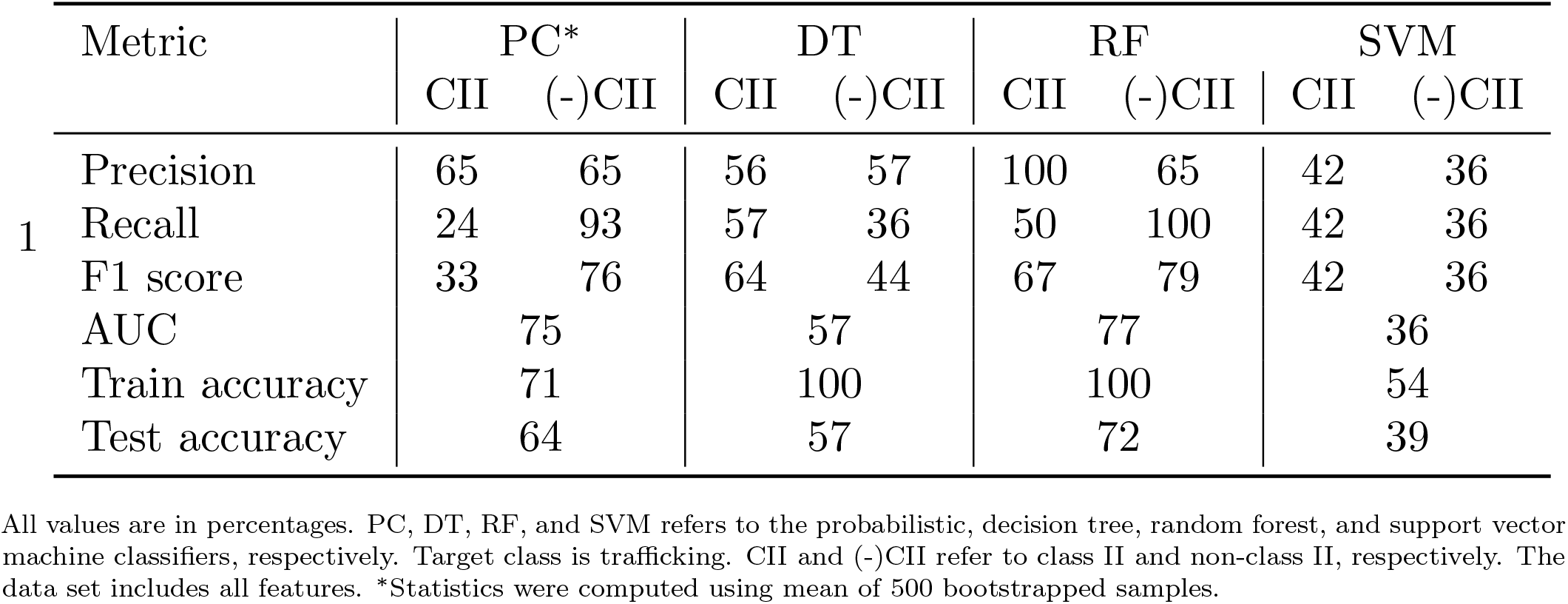
Comparison of DT and RF performance for classifying PAS variants using all features.

| Metric | PC* |  | DT |  | RF |  | SVM |  |
| --- | --- | --- | --- | --- | --- | --- | --- | --- |
|  | CII | (-)CII | CII | (-)CII | CII | (-)CII | CII | (-)CII |
| 1 Precision | 65 | 65 | 56 | 57 | 100 | 65 | 42 | 36 |
| Recall | 24 | 93 | 57 | 36 | 50 | 100 | 42 | 36 |
| F1 score | 33 | 76 | 64 | 44 | 67 | 79 | 42 | 36 |
| AUC |  | 75 |  | 57 |  | 77 |  | 36 |
| Train accuracy |  | 71 |  | 100 |  | 100 |  | 54 |
| Test accuracy |  | 64 |  | 57 |  | 72 |  | 39 |

All values are in percentages. PC, DT, RF, and SVM refers to the probabilistic, decision tree, random forest, and support vector machine classifiers, respectively. Target class is trafficking. CII and (-)CII refer to class II and non-class II, respectively. The data set includes all features. \*Statistics were computed using mean of 500 bootstrapped samples.

#### 3.3.2 Decision tree classifier

We next considered ML approaches that rely on means of classifying the data that are not reliant on probabilities. This is motivated by our observation that several of the classifiers in Fig. S7A are non-monotonic with respect to threshold. We first evaluated DTs, which are well suited for data sets that have a small number of training data points [decision2020]. DTs hierarchically partitions the data by dividing a dataset within a given node according to a threshold value of a feature, such that the resulting partition minimizes the entropy in each ‘leaf’. As an example, we provide in Fig. S16 a DT that partitions the data based on FoldX scores first, followed by hydrophobicity and conservation scores. The first node in this tree divides 52 samples based on whether a variant has a FoldX score above or below 3.77, yielding two leaves with 5 and 13 non-trafficking variants, respectively, as well as entropies of 0.6 and 0.85, respectively. A perfect classifier would partition the variants into two leaves, each with either all CII variants or (-)CII variants, yielding an entropy of zero. Based on this partitioning, the decision tree yields a TPR of 1.0 for the training set versus 0.57 for the test set.

We report in Table 1 F1 scores, recall, precision, accuracy, and AUC (i.e., derived from the ROC curve) assess the DT classification performance. The decision tree using all features yielded a precision of 56% and 57% for the CII and (-)CII variants, respectively. Meanwhile, recalls of 57% and 36% were estimated for the two classes. Overall, this suggests that the decision tree true positive prediction rate is fair and not too reliable for variants with a potential LOF phenotype. The recall is reduced somewhat for traffickers, which suggests a insufficient classification of positive hits. To balance precision and recall, F1 score is reported. Overall, these two factors yield F1 scores of 64% and 44% respectively, which represent harmonic averages of precision and recall. This suggests that the DT classifier does not outperform the probabilistic classifier.

We also computed Receiver operating characteristic (ROC) curves for decision trees trained from three feature sets: bioinformatics alone, MD alone, and all features (Fig. S9). All data sets yield AUCs greater than 0.5, which suggests the classifiers perform better than random. The classifier with the entire feature set yields an AUC of 0.57, which is comparable to that of the MD-only trained classifier. The classifier trained with the bioinformatics features was more accurate (AUC=0.62), which would seem to suggest that the bioinformatics criteria are the most informative for classification. The same was also observed when we evaluate which features contribute most to the DT classification scheme (feature importances). Namely, FoldX has the most significant contribution, followed by RMSF115TO120 and Waters.

#### 3.3.3 Random forest classifier

We next considered an ‘ensembling method’, RFs, which bootstrap and aggregate thousands of randomized decision trees. The bootstrapping and aggregation approach is generally found to improve classification performance and accuracy [yiu2021rf]. RF yielded better results than the DT as reported Table 1. Namely, 100% training and 72% testing accuracy were achieved, with improved precision, recall, and F1 scores for the CII and (-)CII variants over the DT classifier. ROC curve AUCs were also greatly improved for the RF relative to the DT, with values of 0.71, 0.84, and 0.77 for the MD-features alone, bioinformatics features alone and the MD+bioinformatics feature set, respectively (see Fig. S11). The FoldX scores predominated overall all features (0.15), followed by the MD-based features including APBS, 3-10, and Turns (0.10, 0.09, and 0.08).

#### 3.3.4 Support vector machine classifier

Lastly, we evaluated SVMs, which map the input data into high-dimensional feature spaces that are successively partitioned into classes by hyperplanes. These partitions are typically based on polynomial functions [cort1995]. After training SVM models with various feature sets, we report a poorer test accuracy of 39% relative to the DTs and RF approaches, with also a considerably reduced training accuracy (54%) (Table 1). Importantly, both precision and recall were also reduced for the SVM relative to the tree-based models, which resulted in F1 scores of 42 and 36 for the CII and (-)CII variants, respectively. In sharp contrast to the DT and RF approaches, the robust SVM performance is mostly attributed to the bioinformatics-based features. This is shown by the ROC curves in Fig. S13 that were computed for the SVMs trained from MD-only, bioinformatics-only, and combined feature sets, which yielded AUCs of 0.65, 0.11 and 0.35, respectively. Accordingly, we found that the Beta feature importance was the highest among the features used for training, followed by Turns, Hydrophobicity and other MD-related features.

#### 3.3.5 Classifier performance with feature sets

ROC curves showed that RF has better performance than DT and SVM regardless of the feature sets (Fig. 4). We further examined the performance of these classifiers based on the F1 scores. Analyses suggest that RF yielded the best balance between precision and recall when both MD-and bioinformatics-derived features are used. This is demonstrated by the F1 scores shown in Fig. 5 that are marginally superior for the RF CII and (-)CII variants relative to the other models we considered (see solid green and hashed green bars). Additionally, regardless of the model, the F1 scores for CII variants are modestly lower than those for the (-)CII ones, albeit the difference is smallest for the SVM (36 versus 42). These scores also demonstrate that a combined feature set comprising MD-and bioinformatics-based metrics yields the best performance for RF classifications, whereas for DT and SVM, bioinformatics feature set is a major contribution. Interestingly, models trained with the MD-only feature sets performed most poorly, especially for the DT models applied to both CII and (-)CII variants. Altogether, these data suggest that the ML algorithms have better performance relative to a probabilistic classifier and that RF trained from the MD+bioinformatics features yield the highest F1 scores.

**Figure 4:**
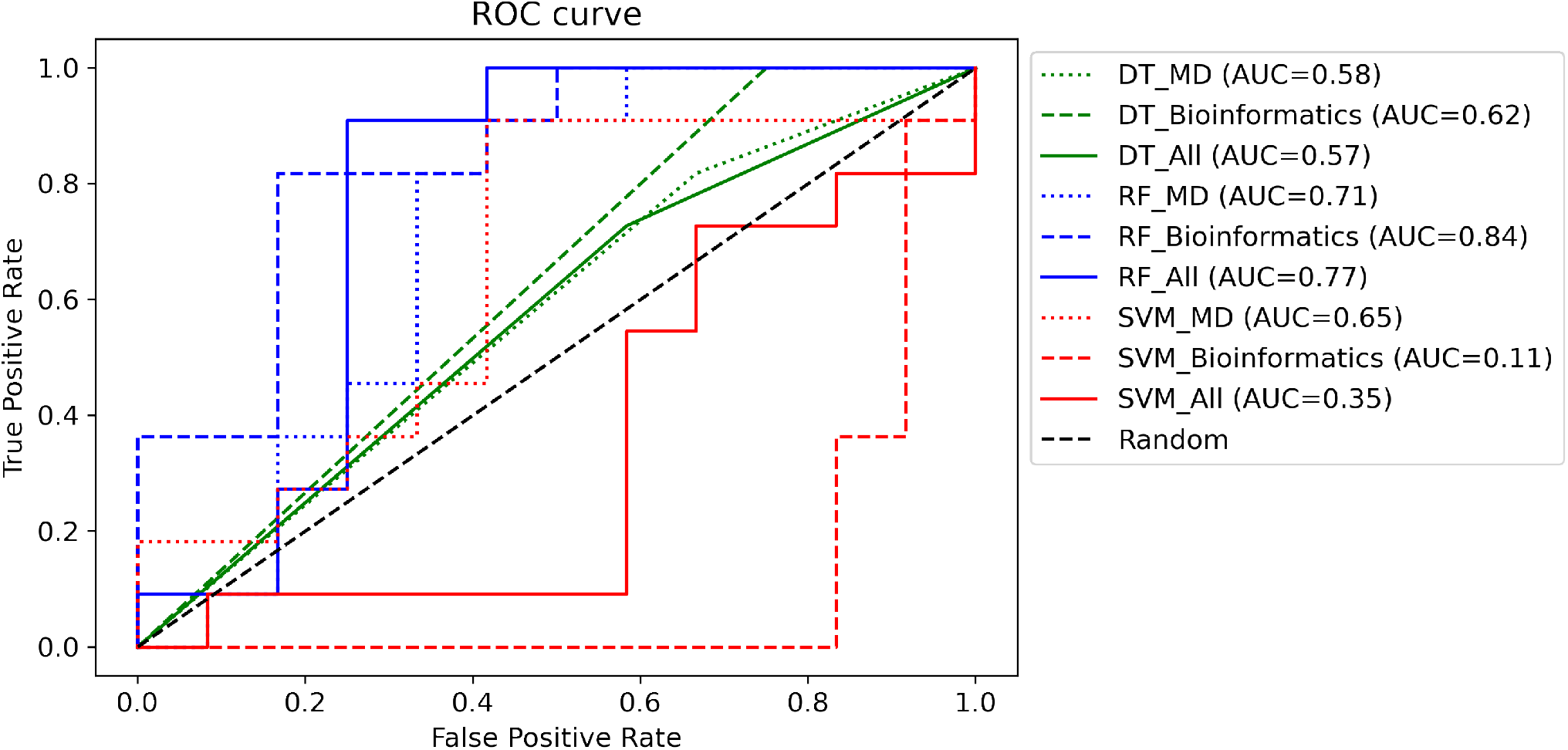
Performance of the DT, RF, and SVM classsifiers on all, MD, and bioinformatics features. ROC curves were used to evaluate predictions on the three data sets. The dotted line represent a random model. For DT and RF, All three sets of features provided good classification (AUC *>* 0.5) with the bioinformatics features being more informative relative to the MD features. In the case of SVM, the MD features yielded superior performance to the bioinformatics and all feature sets.

**Figure 5:**
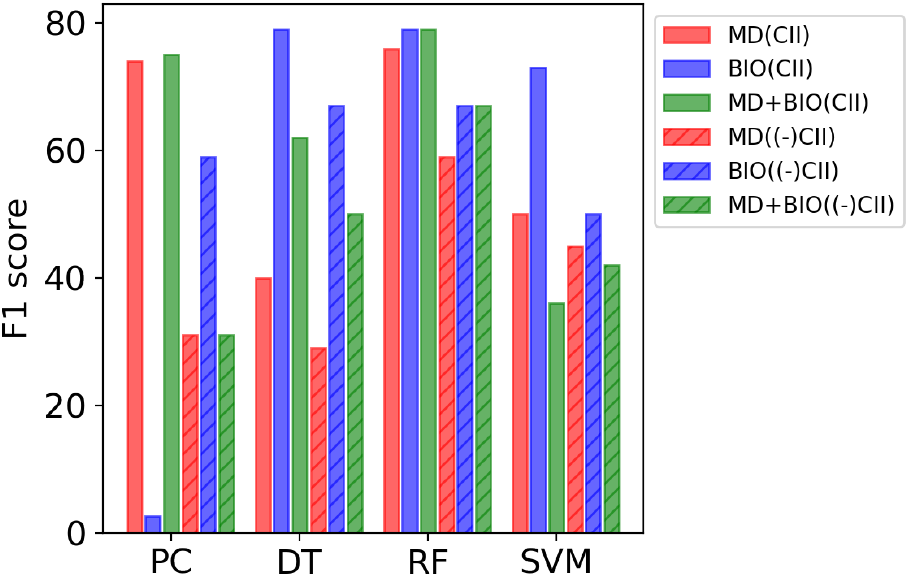
Performances of the classifiers on the three features sets based on the F1 score. MD, bioinformatics, and all-features datasets are colored red, blue, and green, respectively. F1 score is the harmonic mean of recall and precision. CII and (-)CII refer to class II and non-class II, and are labeled as solid and hashed bars, respectively. The RF model overall has supeiror performance over the other classfiers.

Our analyses suggest that the ML classifiers demonstrate the strongest performance (by F1 score) when a full MD and bioinformatics feature set was used. Based on our review of the feature importances we reported for each model (for example, panel B in Fig. S9), conservation score, FoldX energies, total number of waters around the protein, the total number of intra-domain hydrogen bonds and turns were consistently identified. We therefore retrained and tested the classifiers using those features alone to determine if a reduced feature set could improve the models’ performance. Indeed, we found that the F1 scores for CII and (-)CII variants generally improved. Additionally, the ROC performance for each model was also found to improve (DT: 0.57 vs. 0.75, RF: 0.77 vs 0.88, SVM: 0.35 vs. 0.89) (see Fig. S14). Overall, our analyses suggest that by using a pared feature set, the classifier performance is further improved and results in comparable performance by the RF and SVM models.

### 3.4 Application to *de novo* variants

The goal of our study is to provide a framework for predicting a biophysical property in the Kv11.1 channel PASD that correlates with *KCNH2* LOF. Therefore, we applied our approach to several *de novo* variants that have been characterized by our partnering clinic, the Geisinger Medical Center. Those three include N33T, V41I, and V115M, of which all but V41I traffic normally. These variants are summarized in Table 2. We first used the trained ML techniques to classify the variants as soluble versus insoluble using the bioinformatics-based features described above. Given the similar performance among the models we trained, we defined a consensus score that reflects the class predicted by the majority of the DT, RF, and SVM models. Among the variants we considered, R35L, C39R, V41I, D46N, R76H, L87R, F98L, V131A were predicted to be CII variants by consensus, whereas S26C, S26G, R27L, N33T, D46E, C49S, R76L, A79T, R92H, R92C, L107P, V115M, N117S, D119H, M124I, N128S, V131L, M133T were predicted to be (-)CII. We next simulated all of these variants via all atom MD in order to generate their respective MD-predicted features. We subsequently evaluated the variants based on both the bioinformatics and MD-derived feature sets and report the corresponding consensus scores in Table 2. First, the variants N33T, V41I, and V115M characterized by *in vitro* assays were predicted correctly. Interestingly, we found that in general, the classifications predicted (-)CII for nearly all variants except V41I. This was surprising, given that several of these variants were initially classified as CII by the bioinformatics scores alone; if the reclassifications are correct, this would indicate that MD-derived features improve predictions.

**Table 2:** Classification of *de novo* variants predicted by the RF model. Bold-faced entries represent cases that yielded different predictions for bioinformatics-only relative to bioinformatics + MD features.

| Variant | RF Predictions<br>(Bioinformatics) | RF Predictions<br>(Bioinformatics + MD) | JIV<br>Prediction | QTc from EHR<br>(Patients) |
| --- | --- | --- | --- | --- |
| S26G | (-)CII | (-)CII | | $388.00 \pm 0.00$ (1) |
| R27L | (-)CII | (-)CII | (-)CII | $430.25 \pm 11.75$ (2) |
| N33T | (-)CII | (-)CII | (-)CII | $456.33 \pm 4.67$ (2) |
| R35L | (-)CII | **(-)CII** | CII | $411.00 \pm 0.00$ (1) |
| C39R | (-)CII | **(-)CII** | (-)CII | $416.25 \pm 23.25$ (3) |
| V41I | CII | (-)CII | (-)CII | $412.33 \pm 12.25$ (4) |
| D46E | (-)CII | (-)CII | (-)CII | $465.50 \pm 0.00$ (1) |
| D46N | CII | **CII** | (-)CII | $451.05 \pm 27.17$ (12) |
| C49S | (-)CII | (-)CII | (-)CII |  |
| R76L | (-)CII | (-)CII | (-)CII | $434.48 \pm 16.32$ (7) |
| R76H | (-)CII | **(-)CII** | (-)CII | $465.64 \pm 0.00$ (1) |
| A79T | (-)CII | (-)CII | (-)CII | $392.50 \pm 0.00$ (1) |
| L87R | (-)CII | **(-)CII** | (-)CII |  |
| R92H | (-)CII | (-)CII | (-)CII | $465.67 \pm 0.00$ (1) |
| R92C | (-)CII | | (-)CII | $438.47 \pm 0.00$ (1) |
| R92L |  |  | (-)CII |  |
| F98L | (-)CII | **(-)CII** | (-)CII | $429.40 \pm 0.00$ (1) |
| L107P | (-)CII | (-)CII | | $382.00 \pm 100.04$ (4) |
| V115M | (-)CII | (-)CII | | $433.09 \pm 24.42$ (3) |
| N117S | (-)CII | (-)CII | | $452.00 \pm 0.00$ (1) |
| D119H | (-)CII | (-)CII | | $409.37 \pm 13.85$ (4) |
| V131A | (-)CII | **(-)CII** | | $412.00 \pm 0.00$ (1) |
| V131L | (-)CII | (-)CII | | $444.36 \pm 38.83$ (17) |
| M133T | (-)CII | (-)CII | | $456.00 \pm 0.00$ (1) |

In Table 2 we list the clinical QT interval measurements alongside each variant. We emphasize that our modeling team was blinded to the clinical LQTS classification. Based on rubric developed by the American College of Medical Genetics and Genomics and the Association of Molecular Pathology (ACMG/AMP) [Richards2015; Adler2020], variants with QT intervals longer than 460 and 470 ms for males and females, respectively, were deemed LQTS-causing. This cohort of patients presented QT intervals that were overwhelm-ingly sub-threshold. Two patients with at-threshold intervals had histories of syncope or atrial fibrillation. Altogether, none of the patients in this cohort were diagnosed with LQTS. V41I was the only variant that was predicted to be CII using the RF approach based on the bioinformatics and MD features, which was inconsistent with the sub-threshold QT intervals found in patients harboring this mutation. Aside from this incorrectly predicted variant, the remaining 26/27 variants were correctly predicted as (-)CII. Hence, as far as (-)CII variants are concerned, the ML approaches were generally consistent with the clinical phenotype. Importantly, these data may indicate that even if a *KCNH2* variant exhibits a LOF trafficking phenotype, the clinical phenotype may still resemble normal QT intervals. We additionally accompany these data with recently-available *KCNH2* trafficking data reported by Vandenberg *et al*. [ng2021] in Table 2. These data similarly indicate that nearly all variants are (-)CII, with exception to R35L. The R35L variant was found to not traffic by their data in conflict with our model predictions, although this does not appear to be associated with a LQT phenotype per our QTc measurements.

## 4 Discussion

### 4.1 Summary of Results

Bioinformatics and MD-based features were estimated for nearly 80 PASD variants. The selected variants were annotated as trafficking normal or trafficking deficient by a solubility assay described in Anderson *et al*. [Anderson2020] that is a surrogate for cell-based trafficking assays. While few strong trends between features and trafficking competence were identified, machine learning techniques including random forests and support vector machines in particular performed well in correctly classifying the variants by solubility using several features. We then predicted trafficking competence for a set of PASD variants and compared those predictions against variants for which clinical electronic health record (EHR) data were available. Although the computational team was initially blinded to the clinical data, the algorithm predicted nearly all variants, except V41I, would traffic normally. HER ECG data showed that the QT intervals measured in this cohort were not consistent with a LQTS diagnosis, which was mostly consistent with the model predictions. Persons with the V41I variant had not been diagnosed with LQTS or exhibit any obvious LQTS-related phenotypes in the EHR data. This observation reinforces the idea that molecular-and cellular-level phenotypes must be interpreted alongside other clinical considerations for determining LQTS diagnoses.

The American College of Medical Genetics and Genomics and the Association of Molecular Pathology (ACMG/AMP) developed a predictive rubric to assist in the accurate classification of gene variants in Mendelian-linked disorders like LQT2 as “pathogenic, likely pathogenic, uncertain, likely benign, and benign”. [Richards2015; Adler2020] Genetic variants are segregated into these five categories based on allelic frequency of the variant, *in silico* tools, co-segregation analysis, and functional data. This rubric can be applied to variants *KCNH2* to assist in the early identification and prophylactic treatment of people with LQT2 before they suffer a life-threatening event. [Adler2020] There are, however, several limitations to applying this approach. First, the large number of rare non-synonymous variants in *KCNH2* suggest that allelic frequency alone is not a reliable predictor of pathogenicity. [kapa2009] The use of *in silico* algorithms has several challenges in its standardized application and the simplifying assumptions of the algorithms. [Ghosh2017] Many are designed to predict whether a variant disrupts a protein domain and were not intended to be used to infer functional consequence. Another challenge is co-segregation analysis is usually insufficient because the phenotypic information for people identified with a particular variant is typically limited to a family or a few individuals. The most significant gap is the lack of consistency of the functional studies of suspected LQTS mutations. Although initial moderate to high-throughput functional studies of dozens of LQT2-linked mutations show promising results, much more work is needed. [Vanoye2018; Anderson2014; ng2021]

In this study, we used existing functional data to identify ML approaches that could assist in the identification of *KCNH2* variants that predict an impact on channel structure with a functional consequence (disruption of channel protein trafficking) that is known to cause LQT2. Machine learning approaches have been used to probe small molecule binding to the Kv11.1 channel encoded by the *KCNH2* gene. One of the first studies included using machine learning approaches like DTs to predict Torsade de Pointes (TdP)-causing drugs that block Kv11.1 channel current [gepp2006]. More recently, approaches including deep neural network models have been used to predict abrogation of K^+^ current [zhan2019] and ensemble methods for predicting cardiotoxicity of molecular compounds [Liu2020]. In a related study, six ML techniques such as linear regression, logistic regression, naive Bayes, ridge regression, random forest, and neural networks were used to predict Kv11.1 related cardiotoxicity [lee2019]. Fewer ML-based studies aiming to link *KCNH2* variants to LQTS have been published. In a recent study, nearly 15 variants from the Kv11.1 channel PAS, CNBD, or transmembrane domains were classified by their potential to influence trafficking [OliveiraMendes2021]. In addition to their *in vivo* approaches, the study incorporated structural information from the Wang *et al*. Kv11.1 cryo-EM structure (PDB: 5VA1 [Wang2017]). To our knowledge, our study is the first to leverage an extensive set of *KCNH2* variants localized to the PASD to train classifiers. By using an extensive feature set for tens of PASD variants, the predictive power of the ML models described herein is expected to improve.

ML has been extensively used to mine and interpret MD simulation data. Recent examples of these include supervised ML techniques such as logistic regression, random forest, and multilayer perceptron (MLP) to predict interactions between the SARS-CoV-2 Spike Protein and ACE2 [pavl2021], as well as determine physiochemical factors that promote amyloid beta aggregation [guru2021]. Refinement of computational protein structure prediction using ML+MD has also been reported [heo2020]. With respect to variants that could impact protein function, several studies have used MD and ML to predict drug-resistant variants in *Mycobacterium tuberculosis* that is involved in tuberculosis [jama2020; mugu2021], variants that promote CTX-M9 mediated antibiotic resistance [lata2017], variants in epistatic enzyme that impact its stability and aggregation [li2021], and impacts of variants on protein binding dynamics [babb2020] or inducing the phenotypic alterations [garg2019].

We applied several methods to classify the Kv11.1 channel PASD variants considered in this study. There was a general concurrence for the features deemed important for classification, as was shown in Fig. 5. The decision tree method had an advantage in yielding intuitive partitions for classifying the variants. The RF and SVM approaches were less intuitive but were better performing. The probabilistic method performed poorly compared to the nonparametric method, which could be attributed to the nonmonotonic nature of the features.

Among the features we considered, those based on sequence alone had mixed performance, depending on the classification model used. This may in part be due to the relatively low sequence conservation among PASDs [Anderson2020]. The correlation of trafficking with FoldX suggests that the stability of the folded protein is of paramount importance. The apparent relationship between PASD folding stability and trafficking is supported by several features determined from MD simulation. These features include the scores for RMSF, HBond, SASA, H_2_O, and APBS. An increase in the protein RMSF is commonly interpreted as a destabilization of the normally folded structure, although it should be recognized that unfolded proteins can be entropically favorable. The decrease in hydrogen bonds, however, is likely to both decrease the favorable enthalpy of the protein and contribute to its partial unfolding. The loss of hydrogen bonds accounts for the reduction in helical character while increasing turns and 3-10 character, as well as the decrease in beta sheet character that was compensated for by random coil. These relationships are demonstrated in the scatter plots shown in Fig. S3. Commensurate with this unfolding are increases in the protein’s solvent accessible surface area and thereby the number of hydrating waters.

While it is clear that our simulations of some non-trafficking variants present substantial decreases in the number of hydrogen bonds relative to WT, other non-trafficking variants have intermediate reductions that are comparable to the variants that traffic similarly to WT. When other MD properties are considered, including hydration, the ability to differentiate trafficking from non-trafficking variants improved. It is therefore apparent that several molecular features will need to be determined in order to have sufficient data to discriminate between molecular phenotypes, which was the goal of our simulations. It is also important to note that solubility data by immunoblot versus trafficking status from Anderson *et al*. was continuously distributed, with trafficking-deficient variants being more common at lower solubilities. Therefore, there was no clear threshold between folded and unfolded states that could be used for straightforward classification, which advocates for the ML approaches we considered here.

We also found that our MD-based features were mostly consistent with the interpretations outlined in Anderson *et al*. For instance, they reported lesser solvent accessibility of LQT2-associated variants relative to other positions. Additionally, they indicated that increases in the volume of hydrophobic amino acid variants correlated with decreased solubility, although we did not find a strong association. Our simulations, however, indicated that variants involving proline were generally well-tolerated in the folded structure, which conflicts with their suggestion that prolines impose backbone strain upon folding. We also found that a limited number of variants located at the N-and C-termini of the PASD structure trafficked, which is likely because these termini are unfolded in the normal structure. Hence, the MD simulations provide structure-based rationale for the potential of non-trafficking variants to misfold, as evidenced in particular by changes in secondary structure and hydration.

Despite the apparent correlation between FoldX scores and trafficking, there are likely variants that may not negatively impact folding, but rather impair the Kv11.1 channel protein PASD’s interaction with other proteins involved in its trafficking. As an example, exposure of hydrophobic, solvent-exposed regions of the protein may constitute a recognition signal for chaperones of the endoplasmic reticulum [Tsai2002]. Similarly, disruption of the protein’s normal folding may impact different mechanisms of trafficking quality control. These mechanisms could include auxiliary proteins involved in Kv11.1a’s degradation by the proteasome, those recognizing permissive conformations of Kv11.1 channel protein necessary for its trafficking from the ER, or those mediating retrotranslocation of ER-localized Kv11.1 channel protein to or from the cytosol [Gong2005]. To our knowledge, there is a lack of structural data for evaluating this hypothesis, with exception to a PASD structure complexed with the Kv11.1 channel protein cyclic nucleotide binding domain that was determined from x-ray crystallography [hait2013].

### 4.2 Limitations

We discuss several limitations in our current study that could be addressed to improve the classification accuracy. The Anderson *et al*. study utilized a solubility assay as a surrogate for channel trafficking. Ostensibly, misfolded Kv11.1 variants predisposed the proteins to assemble into insoluble aggregates, as has been observed in other misfolded proteins like amyloid beta [floc2006]. Our study identified molecular features that were likely to predict misfolding, but not the aggregation or solubility of the assembled aggregates owing to the difficulty in modeling the latter processes [redl2014]. Along these lines, experimental verification of misfolded proteins could provide a more direct means to link the MD-based features, such as hydrogen bonding, to the Anderson assay. For instance, melting temperatures and circular dichroism have recently been used to probe the proper assembly of PASD variants [Harley2012]. However, these data were only available for a small subset of the variants we considered in our study. Similarly, our analyses focused on the isolated PASD, whereas the determinants of efficacious trafficking may actually reside in the PASD’s impact on the assembled Kv11.1a channel conformation. Such molecular studies are unfortunately not possible with current modeling approaches. Nonetheless, we believe our classification strategy has identified a common set of molecular features in the PASD that correlate with impaired trafficking of the intact Kv11.1 channel. We were also limited by the predominance of non-trafficking versus trafficking variants in our dataset. To better balance the data set, we considered both benign variants and *KCNH2* variants with correctable trafficking defects as ‘trafficking’ (45), whereas only those with uncorrectable defects were labeled as non-trafficking (30) in our model building. This grouping yielded a more even partitioning of the data set, but likely masked more subtle features in the correctable traffickers that would be predictive of the phenotype.

## 5 Conclusions

In this study, we discuss the implementation of a computational protocol comprising ML techniques to predict the trafficking deficient variants of Kv11.1 PASD. We also tested if the addition of structural features obtained from MD simulations to traditionally used bioinformatics scores/features enhanced the predictive power of the most commonly used ML techniques such as DT and RF. We have simulated 75 variants in the PASD, for which the experimental data (solubilities and trafficking) are available via all atom MD. Overall, we have generated 100 μs of MD data and extracted nine different features from these trajectories. Additionally, we have collected three other features such as conservation scores [GLAS2003], change in hydrophobicity, and FoldX energies [SCHY2005]. We have trained both the ML algorithms on three different sets of data: (1) MD data set, (2) bioinformatics scores, and (3) the combination of MD and bioinformatics scores. Furthermore, we evaluated their predictive power on the remaining test data, and we report that the RF algorithm trained on the MD data set alone offered ≈ 76 % accuracy. Particularly, MD features when used along with the bioinformatics scores help improve the identification of non-trafficking class of variants (59 vs. 67 % accuracy). The features that proved to be most critical for the classification of PAS variants for their trafficking defects are FoldX energies, conservation scores, number of intra-domain hydrogen bonds, and number of waters. To our knowledge, this is the first study incorporating MD-derived features into a ML protocol to predict LOF in Kv11.1 channel protein variants.

At a minimum, our strategy for improving the classification of *KCNH2* variant of unknown significance (VUS) comprises a first step to decreasing the number of variants classified as VUS (and subject to more exhaustive functional testing). Given the generality of the approach, the accommodation of new functional testing results is straightforward and in principle will improve in accuracy as more data are collected. For instance, current state-of-the-art medium-throughput electrophysiology methods are being employed by various groups to study and characterize the variants, which could accelerate the expansion of data sets for model training. In addition, structural data for the Kv11.1 channel protein and its intracellular domains are becoming increasingly available, which could unveil new, undiscovered molecular features that are predictive of protein function and ultimately cellular phenotype. Similarly, future studies may adopt data from proteins unrelated to Kv11.1, given the structural similarity of PASDs in other proteins such as EAG (ether-a-go-go) and ELK (EAG-like K^+^) channels [adai2013], as well as the channels like hyperpolarization-activated, cyclic nucleotide-gated channels [Vaccari1999].

## 6 Computational Methodology

### 6.1 Molecular dynamics (MD) simulations

We have used the crystal structure of the isolated PASD(PDB:4HQA [tang2016]) to set up the MD simulations. CHARMM-GUI web server [JO2007] solution builder was used to build the simulation system. Hydrogens were added and then the protein was placed in a TIP3P [SW•JORG83] rectangular water box size of 69 Å × 69 Å × 69 Å with an edge distance of 15 Å on all sides. 0.15 M KCl was added to the water box to neutralize the system. The total # of ions in the system were 27 potassium and 27 chlorides. The total number of atoms in the system was 30,495. Total 80 variants were simulated via all atom MD.

Each system was energy minimized for 100,000 steps using a conjugate gradient algorithm [REID71], and further equilibrated in the NVT ensemble for 5 ns. Production simulations were conducted under periodic boundary conditions in NPT ensemble without restraints. Each system (i.e., the variants that were used to train as well as to test the ML models) on average was simulated for 1.5 *μs* (3 replicas for each system, 500 ns/replica). Each of the de novo variants were simulated for 6 *μs* (3 replicas for each system, 2 *μs*/replica) All systems were simulated with amber16 [amber16]. A 310 K temperature was maintained using a Langevin thermostat and a 1 atm pressure was maintained by Nose-Hoover Langevin piston method [MART94A; FELL95]. The cut-off for nonbonded interactions was 12 Å and the long-range electrostatics were treated using the particle mesh Ewald (PME) method [Darden1993]. Trajectories were saved every 20 ps. Hydrogen bonds were constrained with the SHAKE algorithm [Ryckaert1977]. All simulations were carried out on a local GPU cluster and XSEDE resources. Amber force field (ff19SB [tian2019ff19sb]) was used for the entire system

### 6.2 Machine learning (ML) methodology

Our ML approach comprises four stages (Fig. 6).

**Figure 6:**
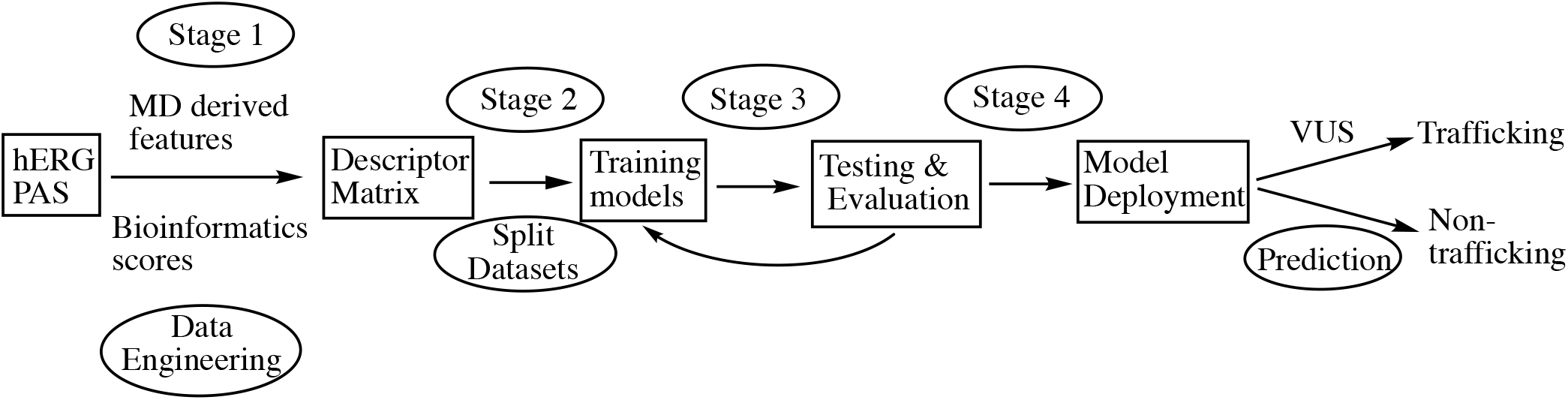
The supervised ML protocol adopted in this study for classifying the variants in the Kv11.1 channel protein. In step 1, the feature set is generated. In step 2, the data are split into training and test sets, and the models are trained on the training data set. In step 3, the models are evaluated on the test data set. The trained model is intended to be deployed for the evaluation of novel VUS to determine their propensity for trafficking.

- **Step 1:** A descriptor matrix is assembled by collecting features for each variant. An excerpt of these data are summarized in Table S1.
- **Step 2:** In this stage, models were trained on the collected features. For this purpose, the data set was split into training (70%) and test (30%) sets, per recommendations in [ROCH2020]. Model training, e.g., optimization of a given model’s hyperparameters, was conducted using the native scikit-learn library.
- **Step 3:** Model performance was assessed on the test data via metrics such as confusion matrix, accuracy, precision, recall, and F1 scores. We emphasized the F1 score because it is a better metric for the imbalanced dataset compared to accuracy [huil2019]. Steps 2 & 3 were carried out in an iterative manner to optimize the performance of the model on the test data.
- **Step 4:** Optimized ML models are deployed to predict the status of novel VUS.

Scikit-learn ML library [pedr2011], numpy and pandas were used for these steps.

#### 6.2.1 Descriptor matrix definition from features

Bioinformatics-based features include FoldX, conservation, and hydrophobicity. The FoldX scores indicate the impact of mutation on domain stability and were computed via FoldX web server [SCHY2005]. The Change in free energy relative to the wild type (ΔΔ*G*) was extracted from the FoldX web server. Conservation scores were estimated via ConSurf server [GLAS2003] to assess the evolutionary rate (conservation) of each residue. Positive and negative values represent high and low conservation, respectively. Conservation score for a residue at a particular position is similar irrespective of the mutation. For example, S26I and S26Y will have similar conservation scores. Hydrophobicity scores of residues were estimated via the Eisenberg and Weiss scale [EISE1982; EISE1984]. Changes in hydrophobicity upon mutation of a residue were estimated compared to the WT. Positive and negative values represent the increase and decrease of hydrophobicity, respectively. Categorical variables (e.g., non-numeric) were converted to numerical values. All variants shown in (Table S1) via all atom MD. Features derived from MD were done using VMD plugins [HUMP96], for which two data points per ns were used. Solvation energy was computed using the Adaptive Poisson-Boltzmann Solver (APBS) at 0.15 M NaCl with solvent and reference dielectric constants of 78.54 and 1.0, respectively ADD REF. A randomized seed was used to split the data into training (70%) and testing data sets (30%). A representative excerpt of the dataset is provided in Table S1. Bootstrapping of the performance metrics described in Section 6.4 used 500 randomized splits of the training and testing data.

### 6.3 Summary of ML and statistical classifiers

ML and statistics-based classifiers were with MD-and bioinformatics-based feature sets. We used the following statistical and machine learning approaches as classifiers for our data.

#### 6.3.1 Conditional probability

We collect *m* features that are normalized as z-scores for *n* variants. The variant *i*’s z-score for its *j*th feature is defined as 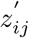. We shift the z-scores for each *j*th feature by the corresponding value for the wild-type structure, 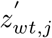, e.g.

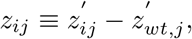

such that *z*_*wt,j*_ = 0 ∀*j* ∈ {*j*_1_, …, *j*_*n*_}. We then define a conditional probability

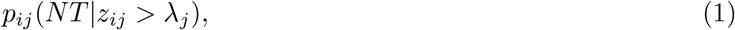

that reflects the likelihood for a variant to be non-trafficking (NT) if its z-score *z*_*ij*_ exceeds some threshold *λ*_*j*_. Eq. 1 is evaluated for all *λ*_*j*_ spanning the minimum and maximum values of *z*_*ij*_ and normalized such that its minimum and maximum are bounded by 0 and 1. We then define a ‘total’ NT probability for the variant *i* as the product of values from Eq. 1 that are determined for each feature *j*

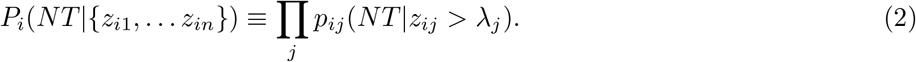

Finally, a variant *i* is deemed non-trafficking if *P*_*i*_(*NT* | {*z*_*i*1_, … *z*_*in*_}) ≥ Λ, otherwise the variant is classified as trafficking. Λ may be tuned to achieve the desired true and false positive rates.

#### 6.3.2 Decision tree

This is a supervised machine learning technique (SML) [soni•supervised•2020], for which the target labels are already known, to classify the variants into trafficking/non-trafficking. We choose this method given that DTs exhibit robust performance [decision2020], are easy to interpret, and that they are nonparametric, meaning that they make no distributional assumptions on the data [tang2017; dt•complete•2019]. Disadvantages include that DTs can be inaccurate, particularly with noisy data, and suffer from overfitting [dt•complete•2019].

We highlight our parameter choices for this model below. Entropy was used as a selection measure [HERS2020]. Hyperparameters for fitting the bioinformatics scores were:criterion=entropy, random state=50, max. depth=None, min. samples leaf=1, max. features=3, and random state=50. ROC curves for individual features were generated by fitting them individually against the target data using the functions provide with SCIKIT-LEARN.

Hyperparameters for fitting the MD models included: criterion=entropy, max depth=None, max.leaf nodes=None, min. samples leaf=1, min. samples split=2, max. features=4, and random state=50.

We observed that when all features were used for splitting at a given node with a decision tree, the DT outputs were biased toward a narrow set of features. Therefore, we have used a maximum of three features (i.e., for the MD dataset), from which one is selected for splitting the data at a given node. The hyperparameters for fitting against all feature data set were: criterion=entropy, max. depth=None, max. leaf nodes=None, min. samples leaf=1,min. samples split=2, max. features=4 and random state=50. These parameters indicate that four were the maximum number of features for splitting at a node.

#### 6.3.3 Random forest

The RF approach is similar to DT, however, instead of fitting the data to a single DT, the models are trained on multiples of DTs, for which each DT is trained on a randomly drawn subset of data. This is repeated 10,000 times, after which the aggregate values of all DTs are estimated [yiu2021rf] The aggregated model is used to classify the test data. In addition to the training data, subsets of features are drawn randomly to fit a given DT. Random forests are a representative ensemble method that has been shown to perform better than the DT when the hyperparameters are appropriately tuned to fit the training data [yiu2021rf]. Other advantages include that they are less prone to overfitting than DT, can work with high dimensionality data efficiently, overcome data points with an incomplete feature set, and identify outliers [yiu2021rf].

The hyperparameters we chose for the RF model when fitting against bioinformatics data include: bootstrap=True, #estimators=10,000, criterion=entropy, max. depth=None, max. features=3, max. leaf nodes=None, min. samples leaf=1, min. samples split=2, random state=50. A maximum of two features were drawn randomly for splitting a node in RF model.

The hyperparameters used to fit RF models with md data include: bootstrap=True, #estimators=10,000, criterion=entropy, max depth=None, max. features=4, max. leaf nodes=None, min. samples leaf=1, min. samples split=2, random state=50. Any tree can only use up to a maximum of three randomly selected features in RF model for node splitting.

Lastly, the hyperparameters for all feature set are: bootstrap=True, #estimators=10,000, criterion=entropy, max depth=None, max. features=4, max. leaf nodes=None, min. samples leaf=1, min. samples split=2, random state=50. Any given tree can only use the maximum four features in RF model while splitting at a node, and these four features are randomly drawn.

#### 6.3.4 Support vector machines

SVM models were constructed similar to the procedures described above. 70% of the data was used to train the classifier and 30% for the testing the trained models. Unlike the others ML methods explained above, SVM generates a hyperplane to separate the classes in the high-dimensional feature space [bzdo2018]. Hyperparamaters used to fit the SVM models (for all three data sets) were: gamma=‘auto’, random state=50, max. iter=10000, kernel=‘linear’, probability=True. A default value of 1 was used for C (also called as the regularization parameter). C and regularization are inversely proportional, i.e., greater the value of C lower is the regularization for a model [bzdo2018]. Gamma is the kernel coefficient and gamma=‘auto’ enables the use of 1/number of features for a given model []. Typically, a large value of gamma leads to large bias and low variance and vice versa. Max. iter = −1 indicates that there is no limit on the maximum iterations.

### 6.4 Model performance assessments

We utilize several metrics to assess model performance. We report the models’ accuracy, which is formally defined as

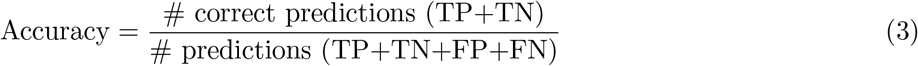

where TP, TN, FP, FN are true positives, true negatives, false positives, and false negatives, respectively. The accuracy provides a simple assessment for the probability of correct positive assignments. However, this metric can yield misleading results, especially for imbalanced datasets [jose2017], therefore we also report the precision. The precision reflects the validity of the model, that is, the percentage of positives that are true positives:

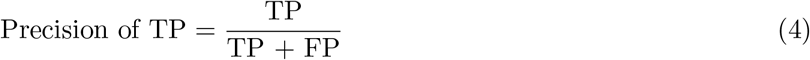

The recall, which is also frequently referred to as sensitivity, indicates the percentage of true positives that were correctly identified:

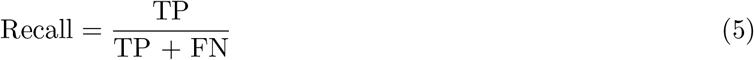

Lastly, we report the F1-score, which is the harmonic average of recall and precision.

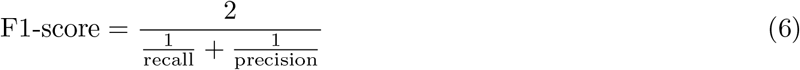

The F1 score measures the balance between the precision and recall of the model and is the de facto standard for comparing ML models.

We also accompany these data with a confusion matrix, which is a N × N matrix used to evaluate the performance of a model/classifier on test data when the true values are known. N is the number of target classes, which typically are four in the case of binary classification (false versus true). Performance was also assessed via ROC curves that report the true positive rate (TPR) and false positive rate (FPR) as a function of threshold The Area under the curve (AUC) for each curve designates the model performance. Namely, an inefficient model for which TPR=FPR will yield an AUC of 0.5 relative to an optimal model, for which AUC = 1.0 (FPR=0% and TPR=100%) All calculations were conducted using numpy and the SCIKIT-LEARN package.

## 7 Acknowledgements

Research reported in this publication release was supported by the American Heart Association under grant number 20IPA35320141.

## S1 Supplementary Information

**Figure S1:**
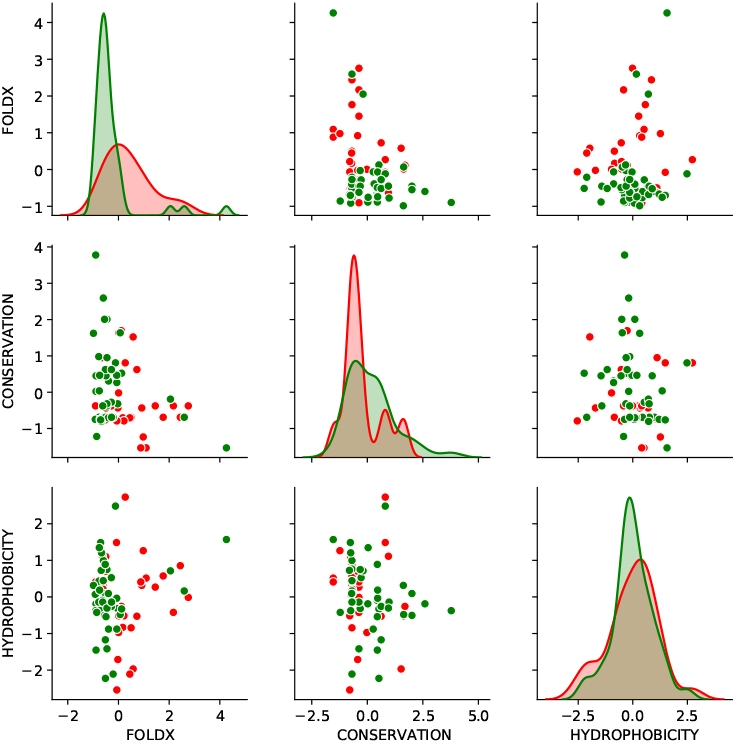
Distribution of the bioinformatics data. Green and red represents trafficking and non-trafficking variants, respectively.

**Figure S2:**
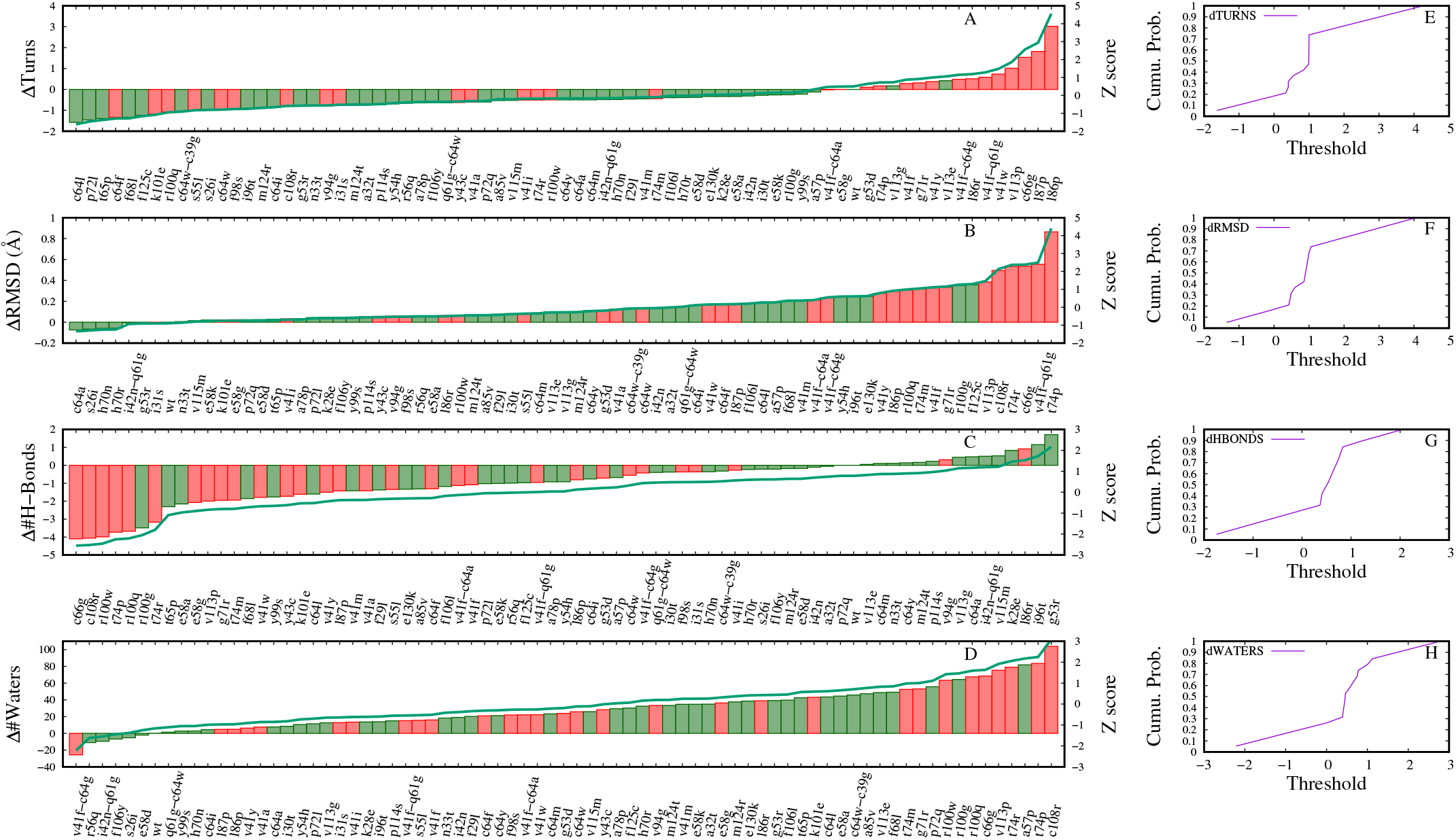
Comparing variant classification by selected MD features. Shown in the left panels are ranking of variants by MD features such as turns, protein backbone RMSD, # H-Bonds, and # waters. On the y-axis, change in respective MD feature value relative to the WT is shown. Waters within 4 Å of the protein were calculated. In the right panels, cumulative probabilities of individual MD features as a function of threshold is shown. See Fig. S7 for conditional probabilities.

**Figure S3:**
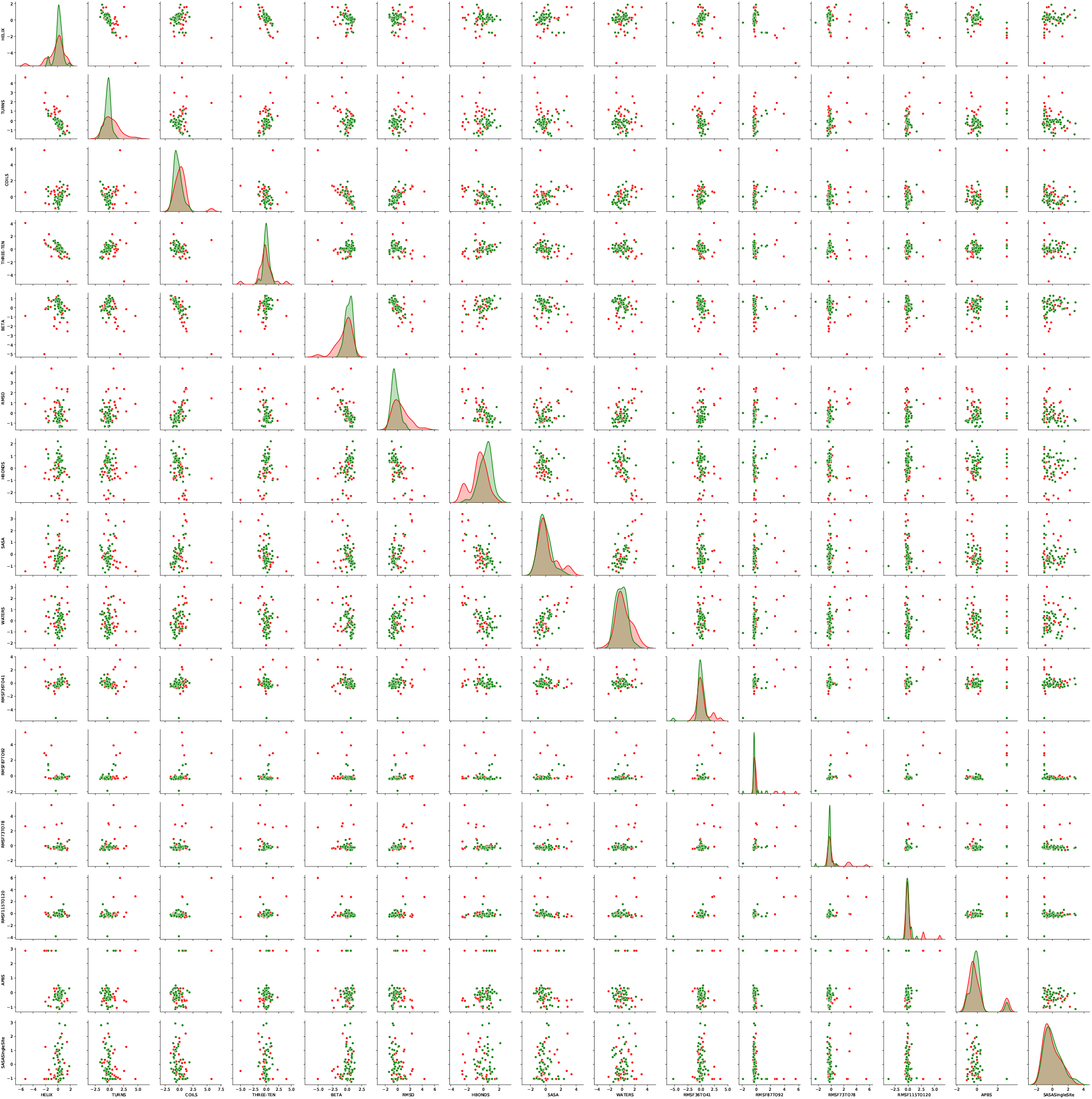
Distribution of the MD data. Green and red represents trafficking and non-trafficking variants, respectively.

**Figure S4:**
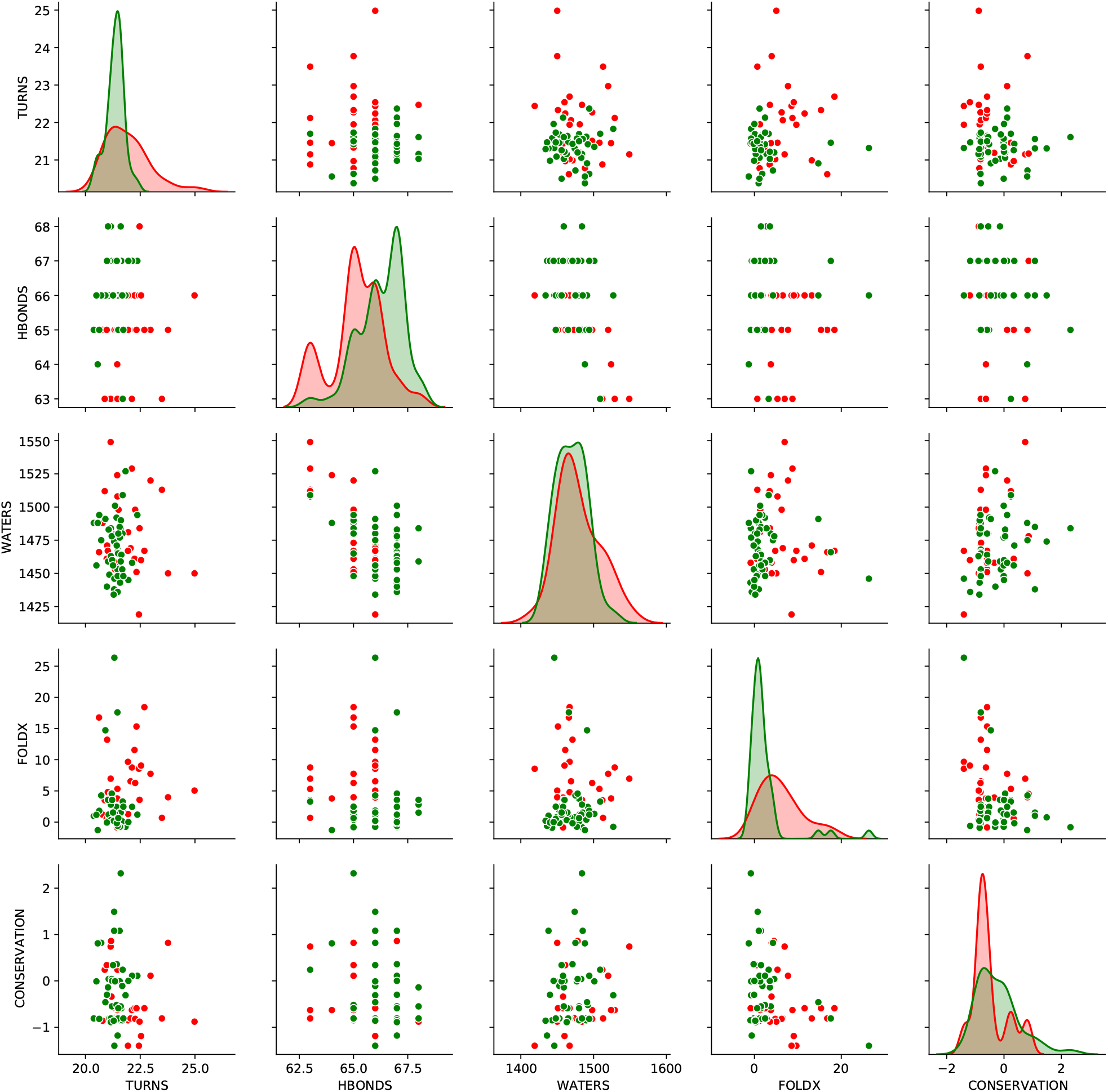
Distribution of the best features. Green and red represents trafficking and non-trafficking variants, respectively.

**Figure S5:**
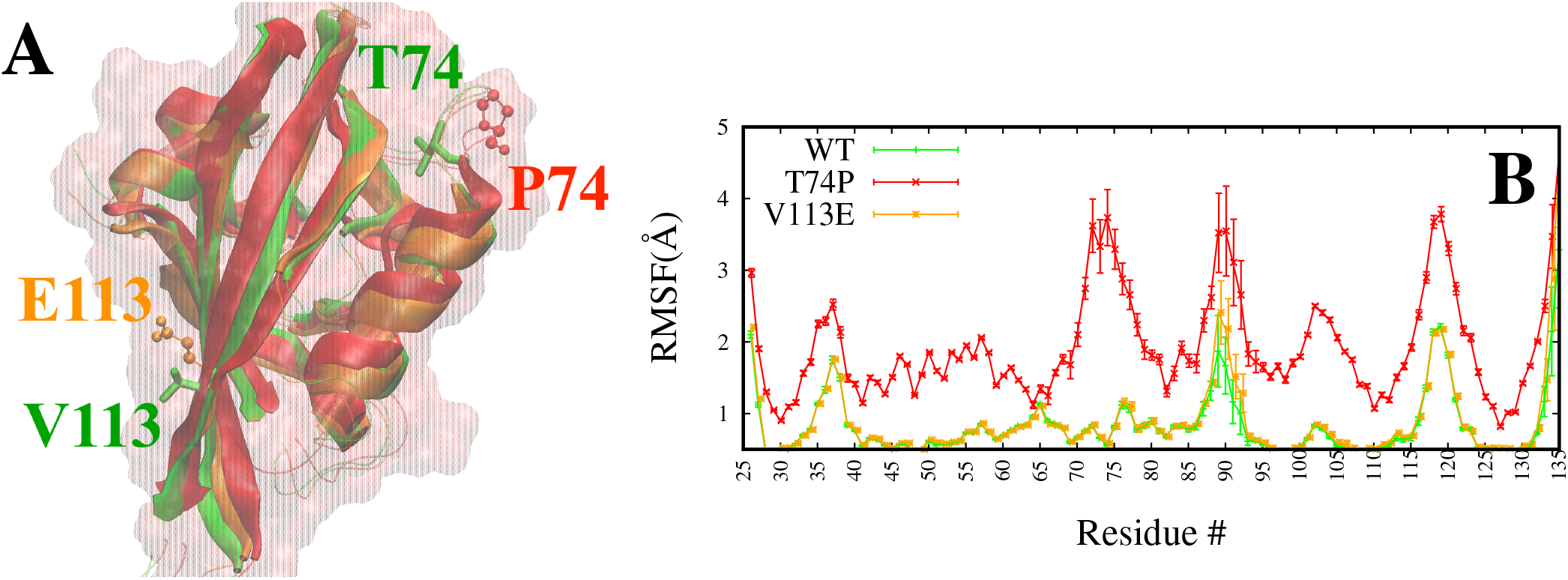
MD structures and RMSF. (A) Superposition of representative MD snapshots of WT (green), V113E (orange), and T74P (red) variants. Residues of WT are shown in the licorice representation and of the others as ball and stick models. T74/P74 is in the loop and E113/V113 is stacked between *β*-strands. (B) Comparison of RMSFs of WT, V113E, and T74P. T74P is pathogenic (non-trafficking) and V113E is benign (trafficking) [Anderson2020].

**Figure S6:**
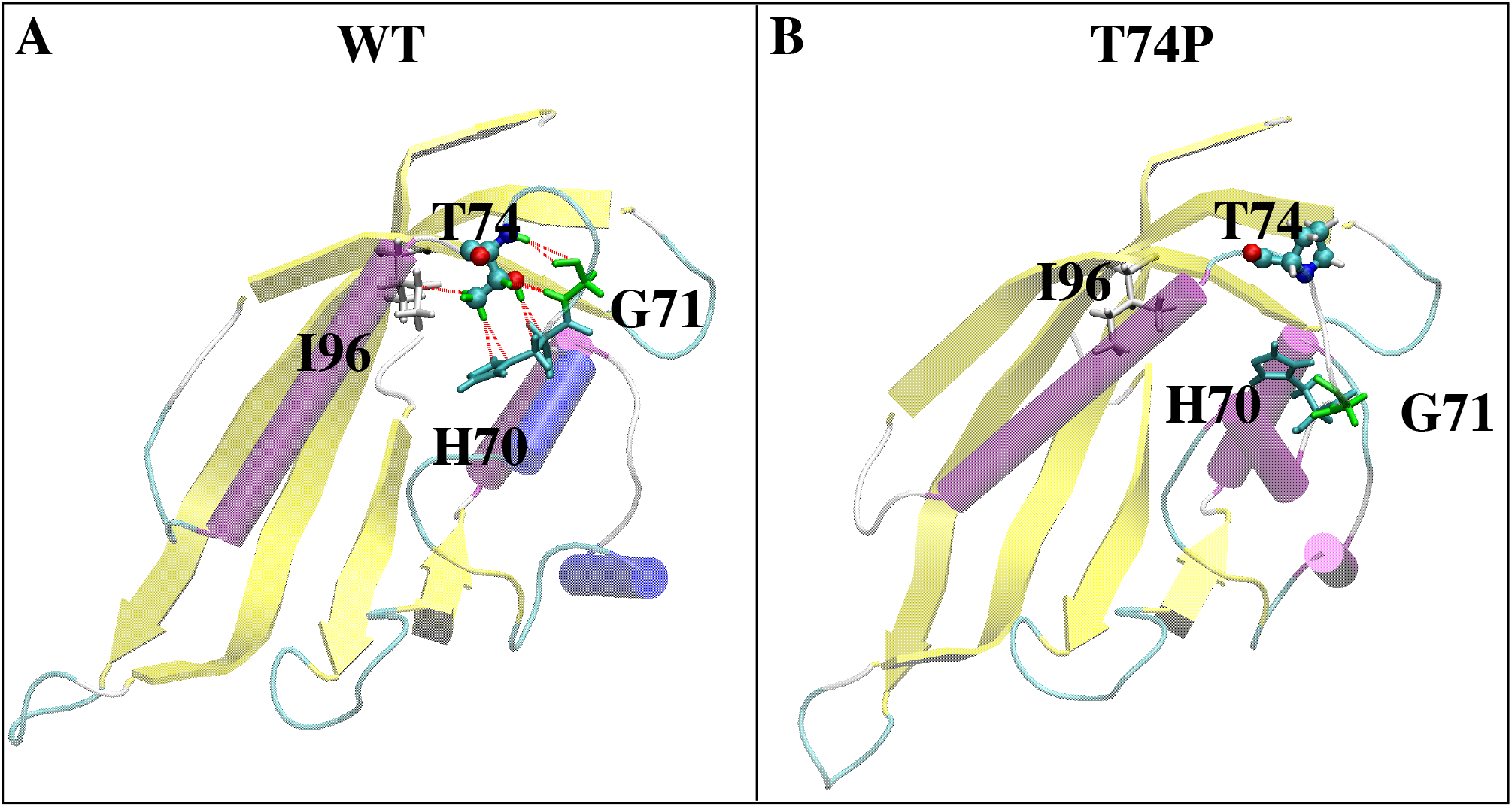
MD representative snap shots showing the hydrogen bond network near the site of mutation in WT (A) and T74P (B) variant. There are at least three hydrogen bonds (red lines) that T74 forms with H70 (cyan), G71 (green), and I96 (silver/white), respectively, in the WT. These hydrogen bonds are absent in the T74P variant.

**Figure S7:**
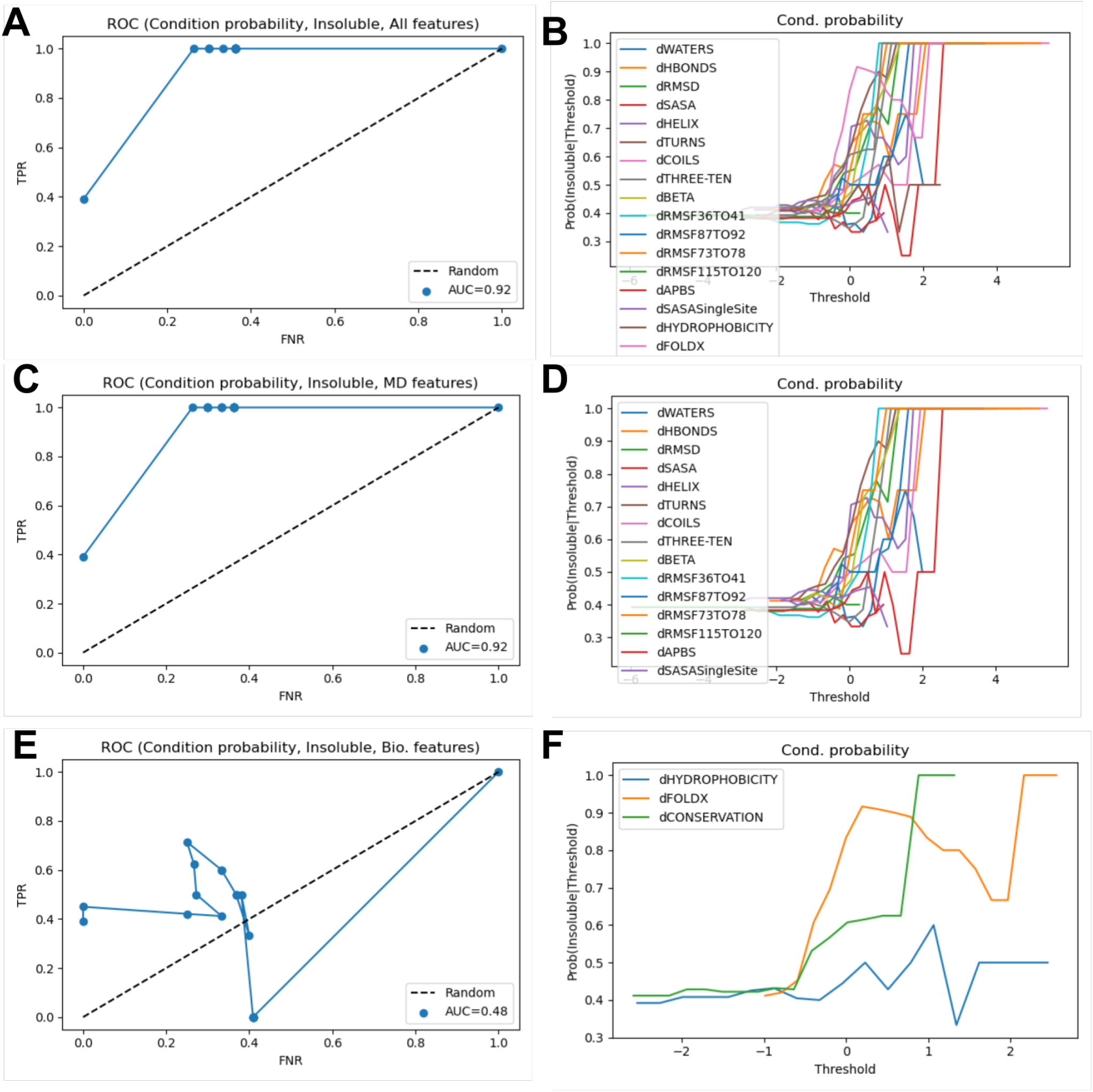
Performance of the probabilistic classsifier on all (A-B), MD (C-D), and bioinformatics features (E-F). Data shown here are extracted from one representative bootstrapped sample. Statistics of the performance of the classifier is reported in Table 1 based on 500 bootstrapped samples. Left column, ROC curves evaluating the performance of the classifier in predicting the insoluble/non-trafficking variants. Dotted lines represent the random model, *i.e*., a model that lacks any predictive capability (AUC = 0.5). Right column, conditional probability as a function of threshold.

**Figure S8:**
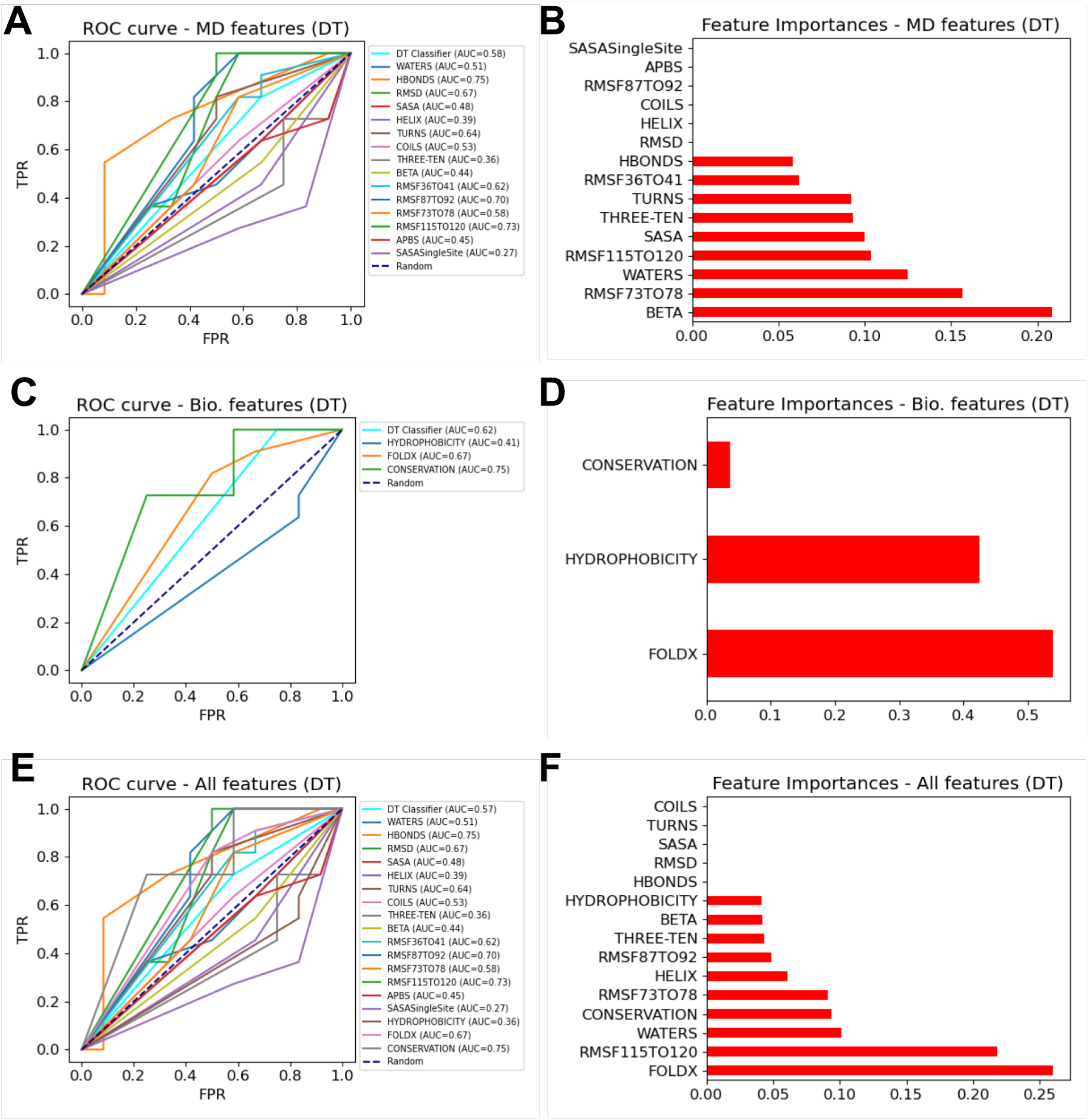
Comparison of DT performance on MD, bioinformatics and all-feature sets. ROC curves estimated for the DT models for all three datasets are shown in the left panels. AUC refers to area under the curve. ROC curves were estimated for the entire feature set as well as for the individual features. Shown in the right panels are feature importances estimated by DT models for the three datasets, respectively.

**Figure S9:**
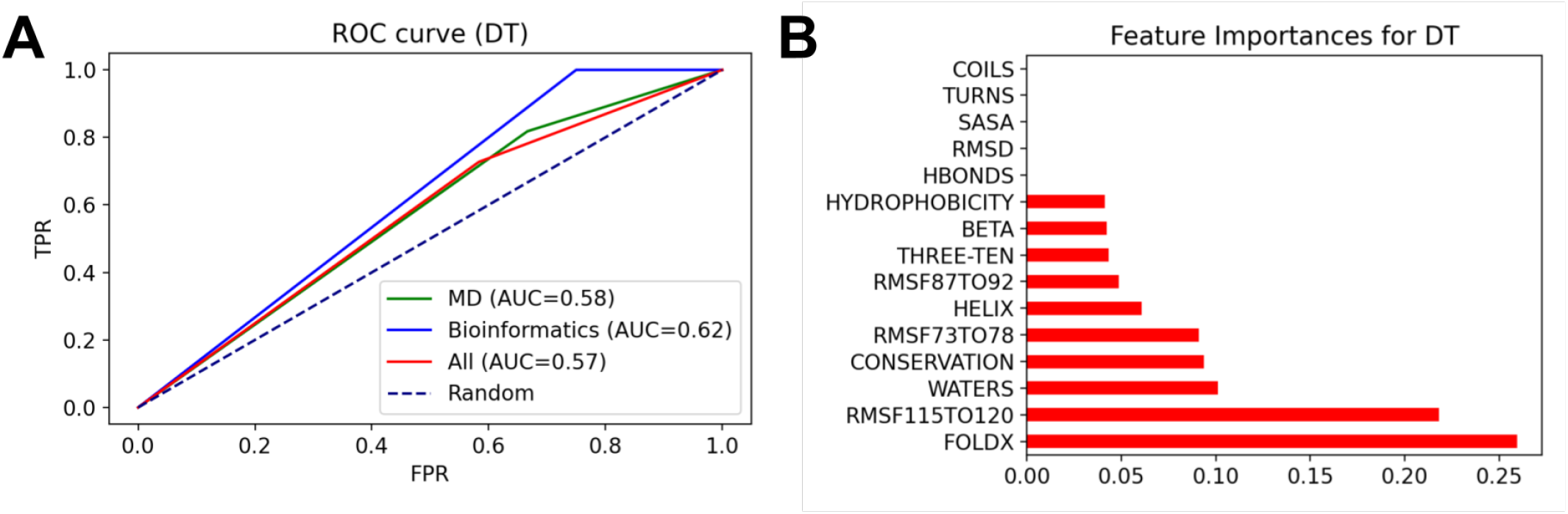
Performance of the DT classsifier on all, MD, and bioinformatics features. (A) ROC curves were used to evaluate DT predictions on the three data sets. The dotted line represent a random model. All three sets of features provided good classification (AUC *>* 0.5) with the bioinformatics features being more informative relative to the MD features. (B) Feature importances estimated via the DT model for the all features data set. It indicates that FoldX, RMSF, and Waters characters contribute most to the classification.

**Figure S10:**
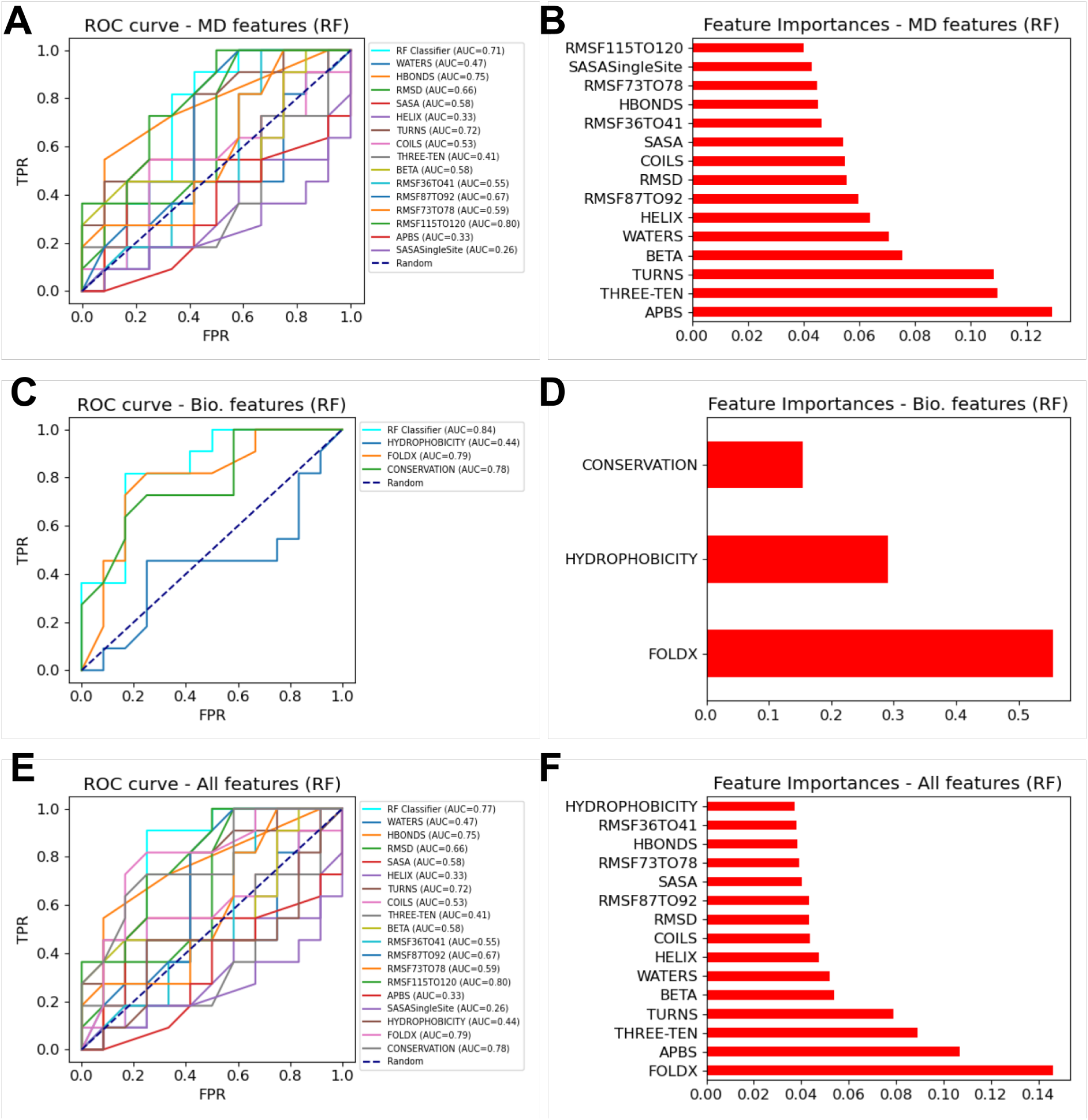
Comparison of RF performance on MD, bioinformatics and all-feature sets. ROC curves estimated for the RF models for all three datasets are shown in the left panels. Shown in the right panels are feature importances estimated by RF models for the three feature sets, respectively.

**Figure S11:**
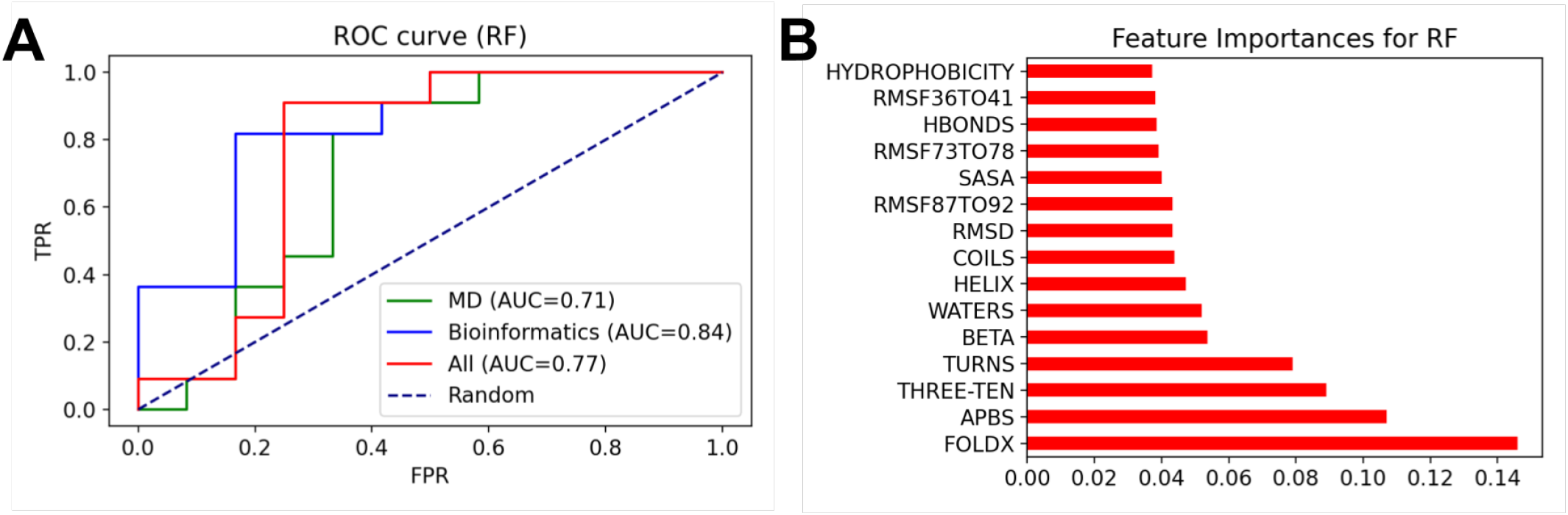
Performance of the RF classsifier on all, MD, and bioinformatics features.(A) ROC curves were used to evaluate DT predictions on the three data sets. (A) The dotted line represent a random model. All three sets of features provided good classification (AUC *>* 0.5) with the bioinformatics features being more informative relative to the MD features. (B) Feature importances estimated via the DT model for the all features data set. It indicates that FoldX, APBS and 3-10 characters contribute most to the classification.

**Figure S12:**
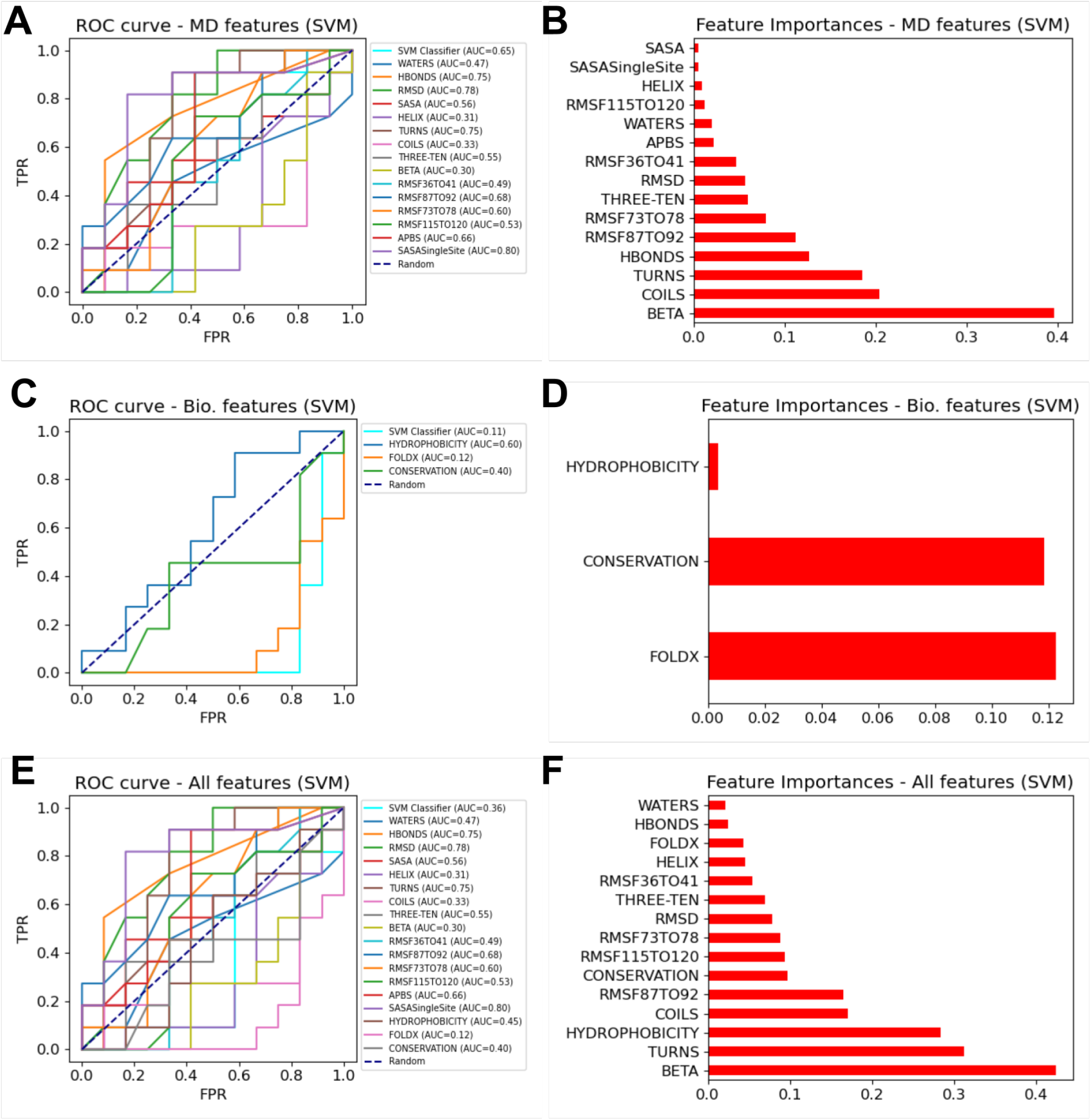
Comparison of SVM performance on MD, bioinformatics and all-feature sets. ROC curves estimated for the SVM models for all three datasets are shown in the left panels. Shown in the right panels are feature importances estimated by SVM models for the three feature sets, respectively.

**Figure S13:**
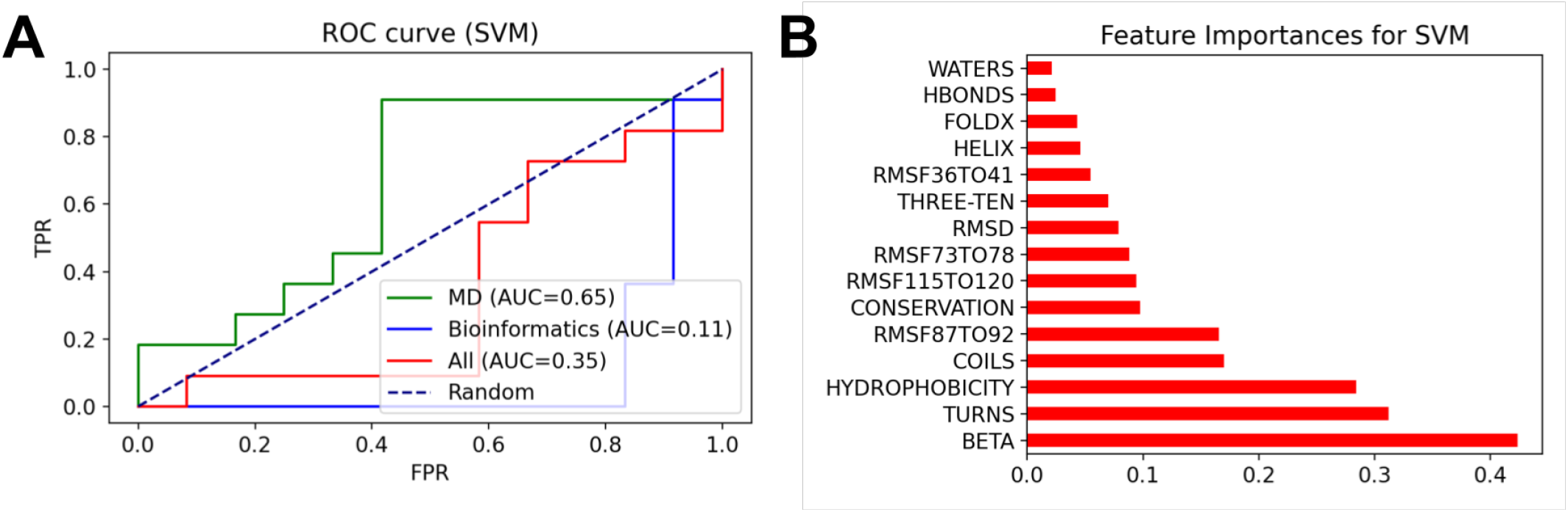
Performance of the SVM classsifier on all, MD, and bioinformatics features.(A) ROC curves were used to evaluate DT predictions on the three data sets. (A) The dotted line represent a random model. Only the MD features provided good classification (AUC *>* 0.5) whereas the bioinformatics features yielded poor performance. (B) Feature importances estimated via the DT model for the all features data set. It indicates that Beta, Turns, and hydrophobicity characters contribute most to the classification.

**Figure S14:**
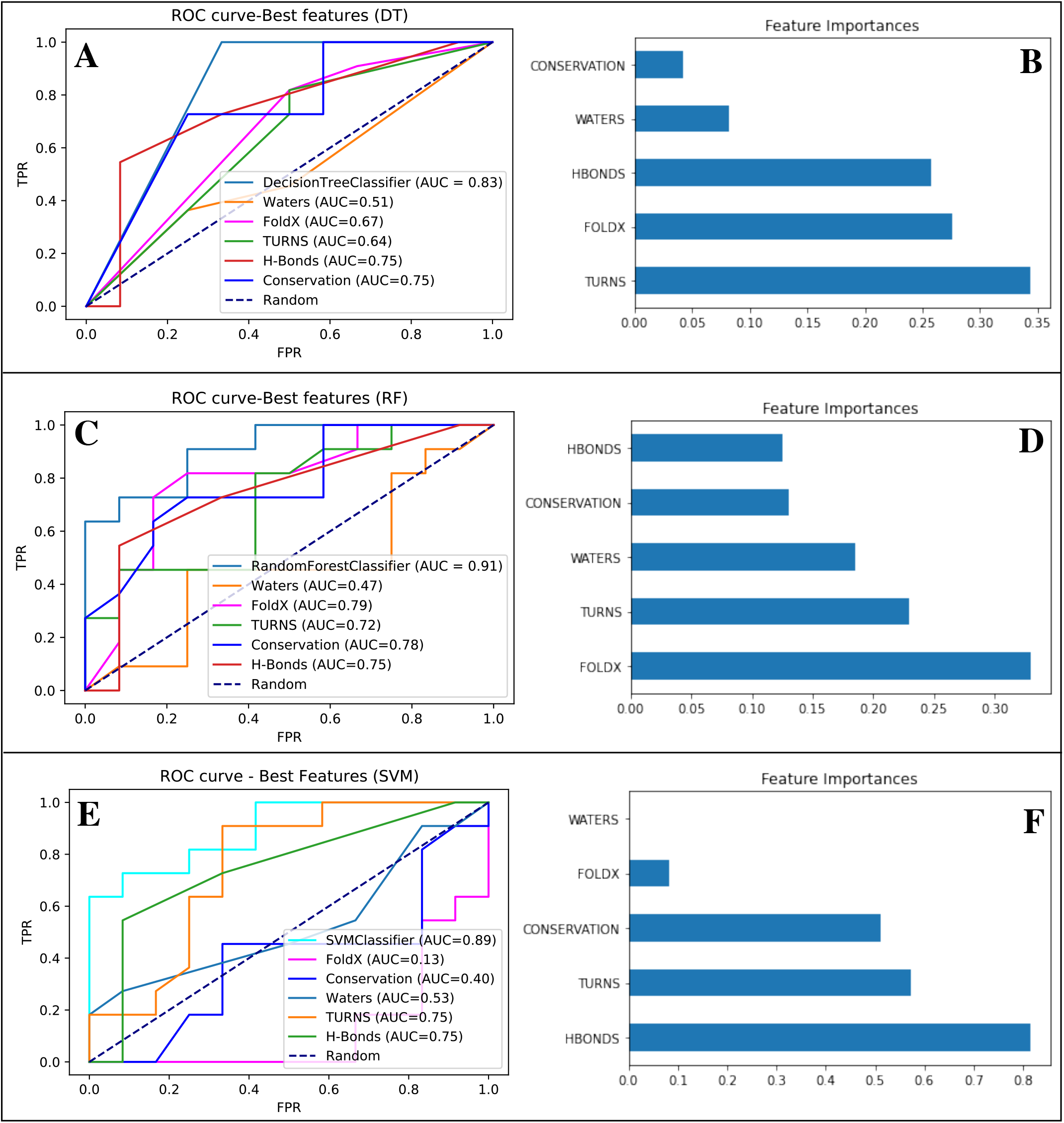
The best five features (i.e., conservation, H-Bonds, waters, FoldX, and turns) were used to train DTs (A,B), RFs (C,D) and SVMs (E,F). For the DT and RF classifiers, maximum two features were allowed to split at a given node. ROC curves for all three models are shown in the left panels. Shown in the right panels are feature importances estimated by all three models, respectively.

**Figure S15:**
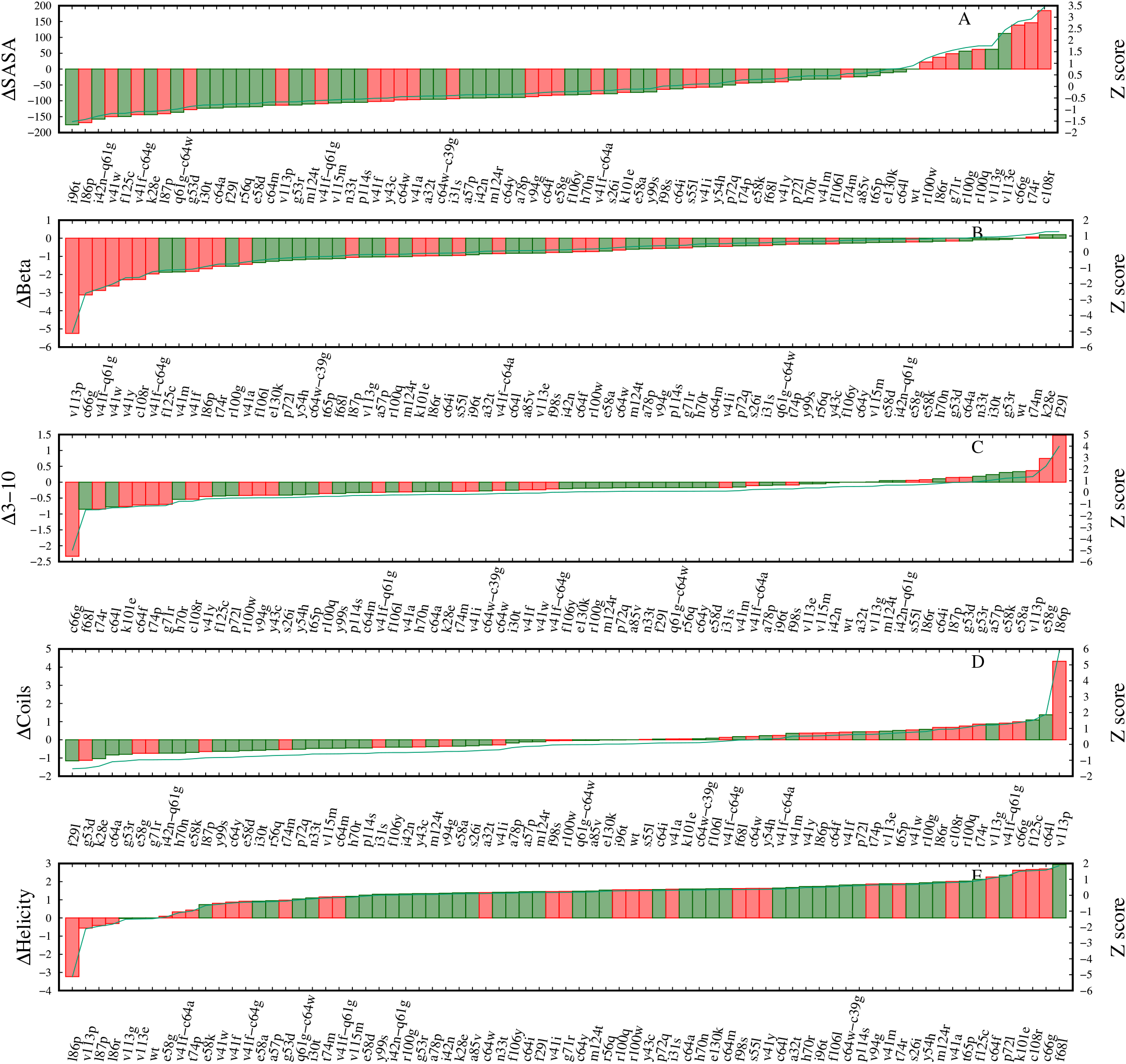
Ranking of variants by select subset of MD features. Red and green represent non-trafficking and trafficking variants, respectively. Shown on y-axis is change in the respective score for a variant compared to the WT. Y2-axis represent z-score.

**Figure S16:**
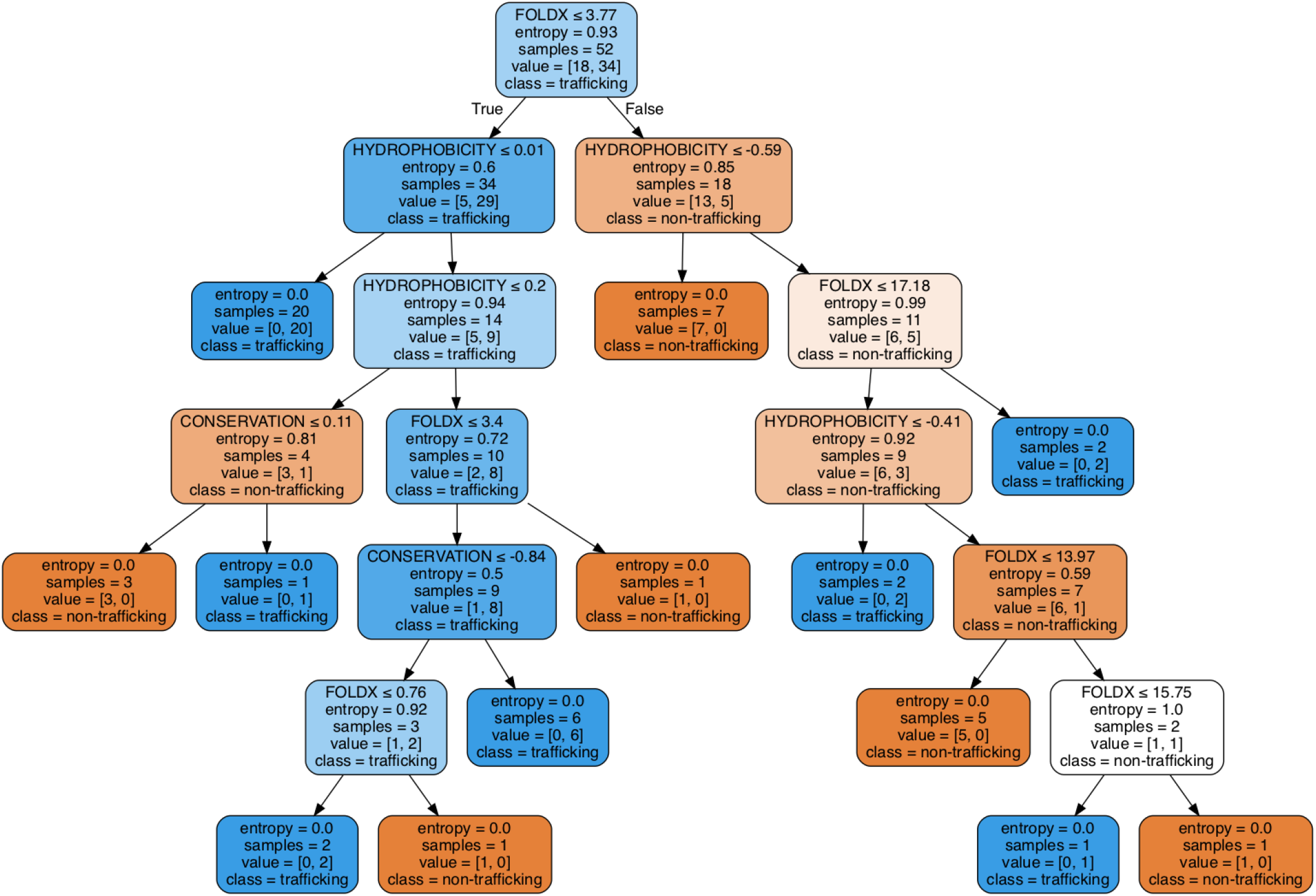
Classification of variants by the DT model when bioinformatics scores were utilized. Orange and blue boxes represent entropies of 0 and 1, respectively. FOLDX score was selected in the root node (i.e., the first node in which data splitting started) to split the dataset. Splitting stops when entropy is zero at a give node (called as a leaf node) and the splitting continues for nodes whose entropies are *>*zero (called as internal nodes).

**Figure S17:**
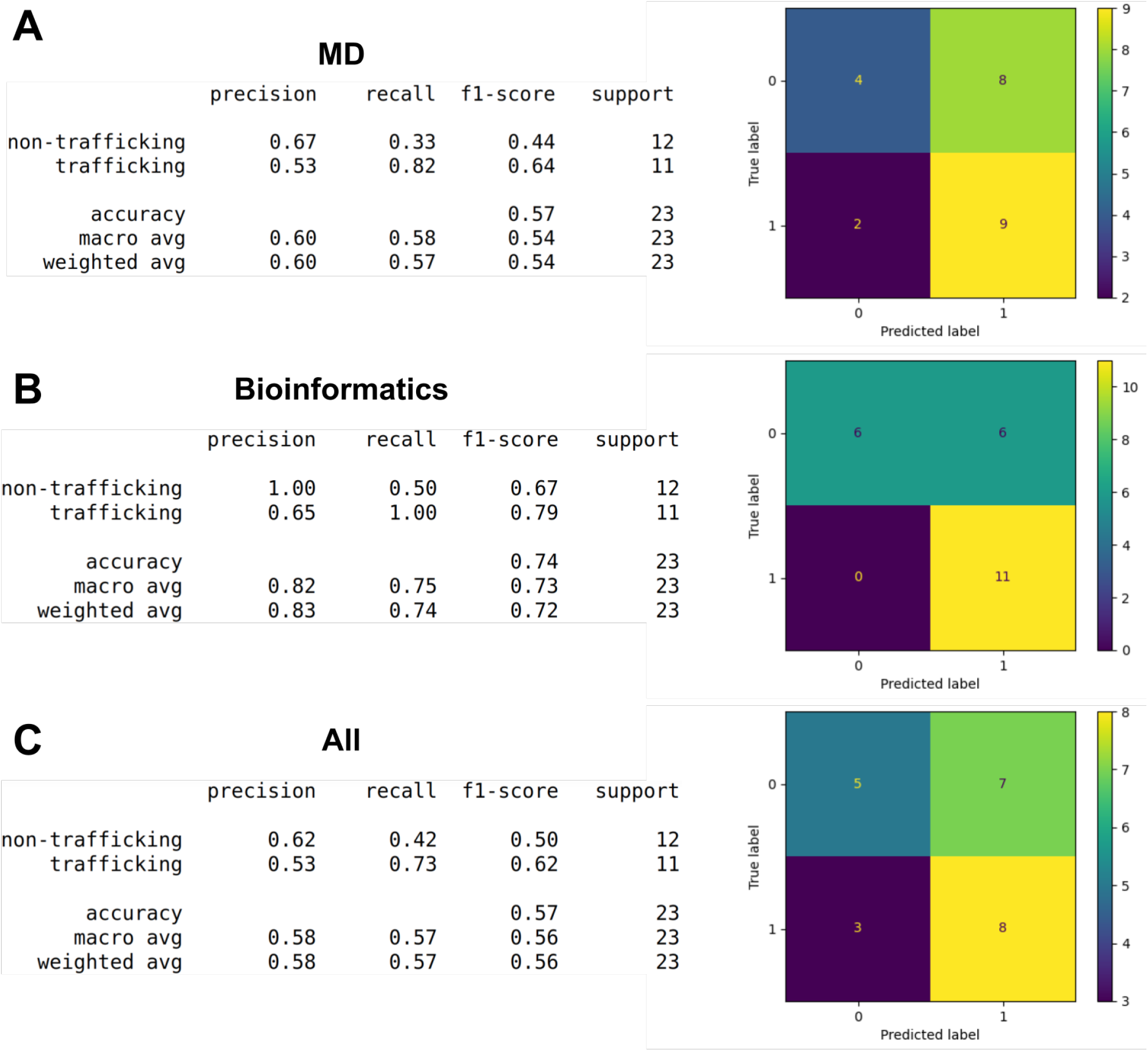
Evaluating the DT model performance on three feature sets (MD, bioinformatics and combination) of PAS domain. Shown on right are the confusion matrices that help evaluate the models performance when the true values of the test data are known. The target classes in the confusion matrices are true positive, true negative, false positive and false negative. Support defines the number of data points in the test data set.

**Figure S18:**
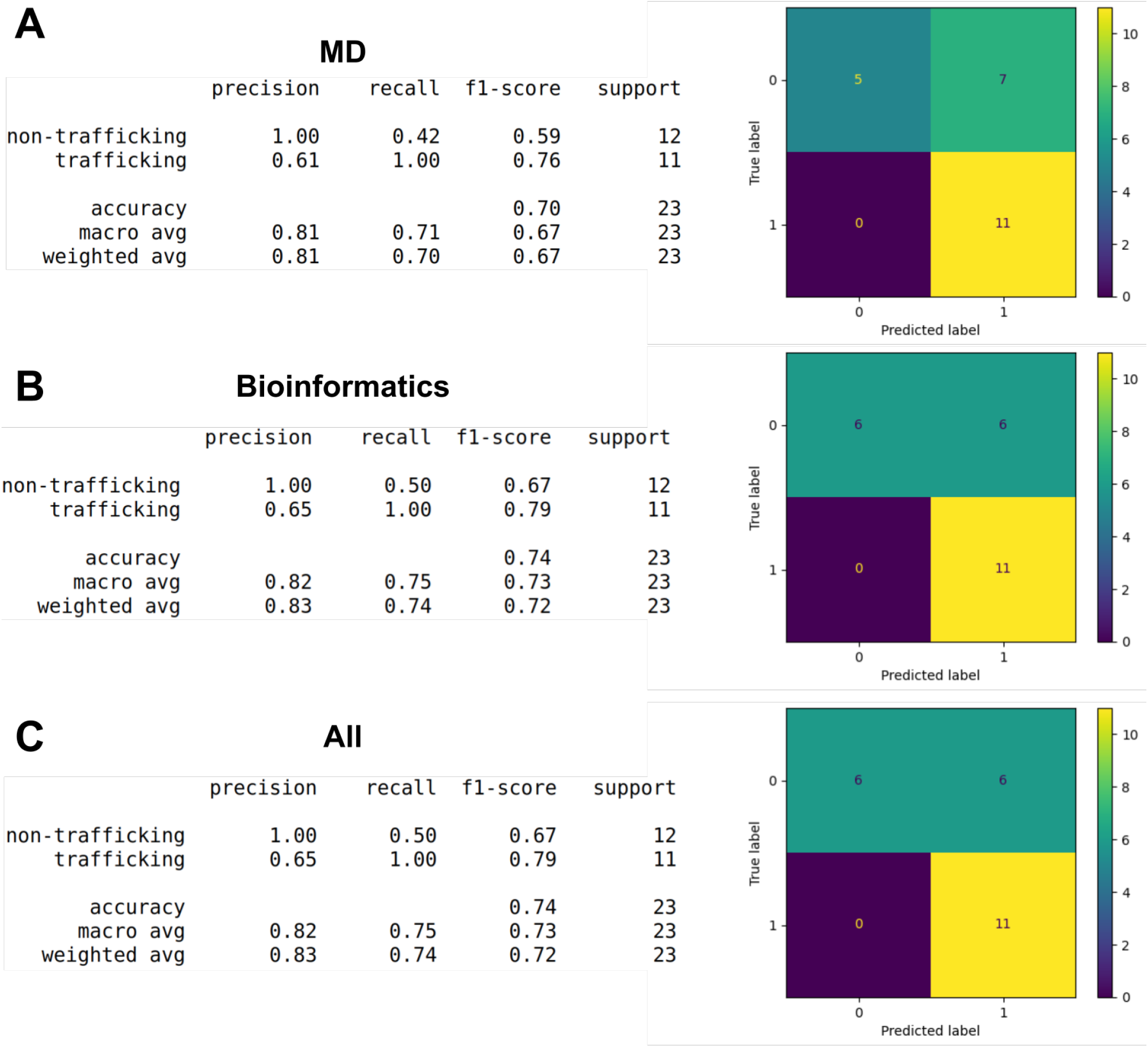
Evaluating the RF model performance on three feature sets (MD, bioinformatics and combination) of PAS domain. Shown on right are the confusion matrices that help evaluate the models performance when the true values of the test data are known. The target classes in the confusion matrices are true positive, true negative, false positive and false negative. Support defines the number of data points in the test data set.

**Figure S19:**
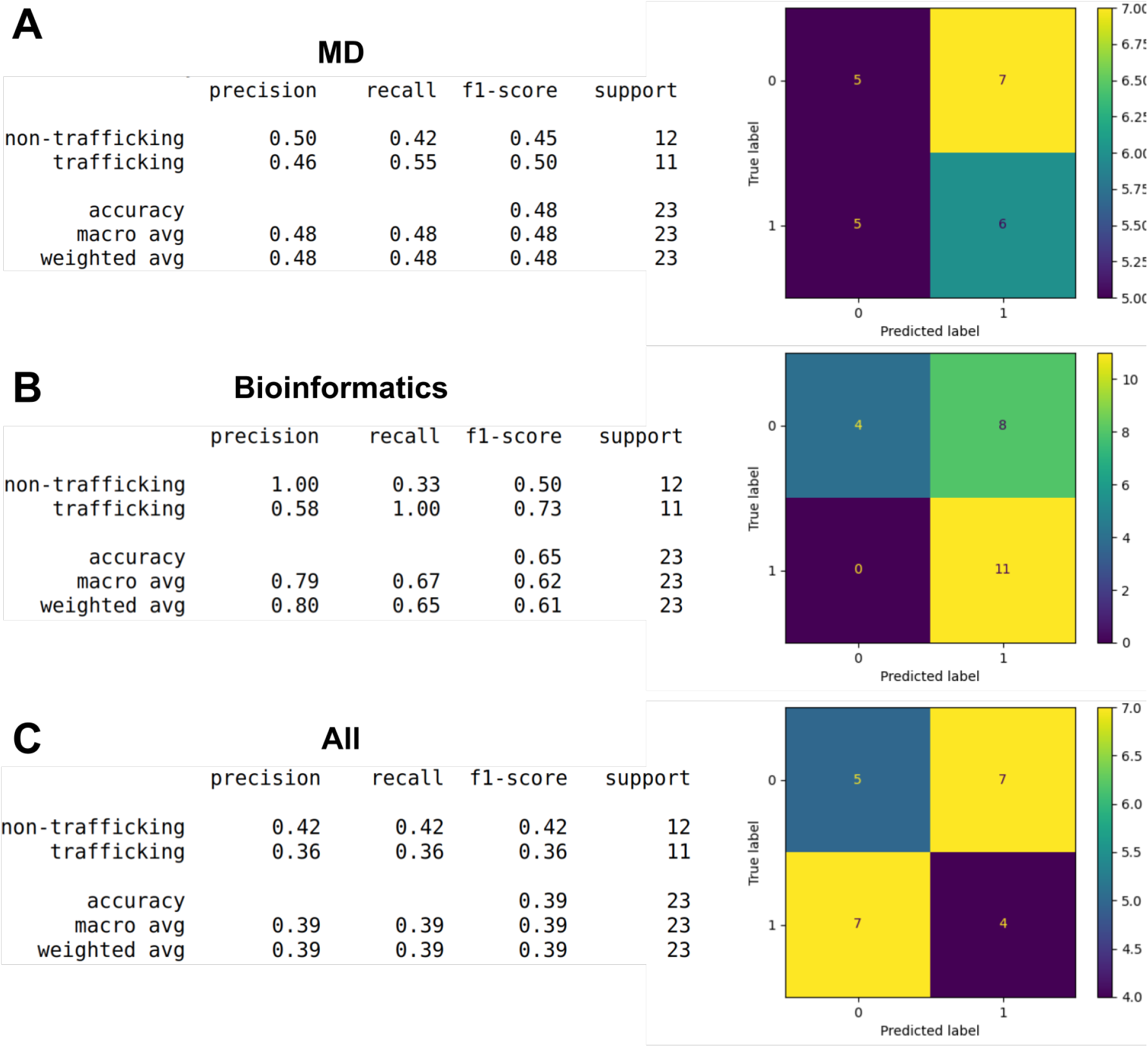
Evaluating the SVM model performance on three feature sets (MD, bioinformatics and combination) of PAS domain. Shown on right are the confusion matrices that help evaluate the models performance when the true values of the test data are known. The target classes in the confusion matrices are true positive, true negative, false positive and false negative. Support defines the number of data points in the test data set.

**Figure S20:**
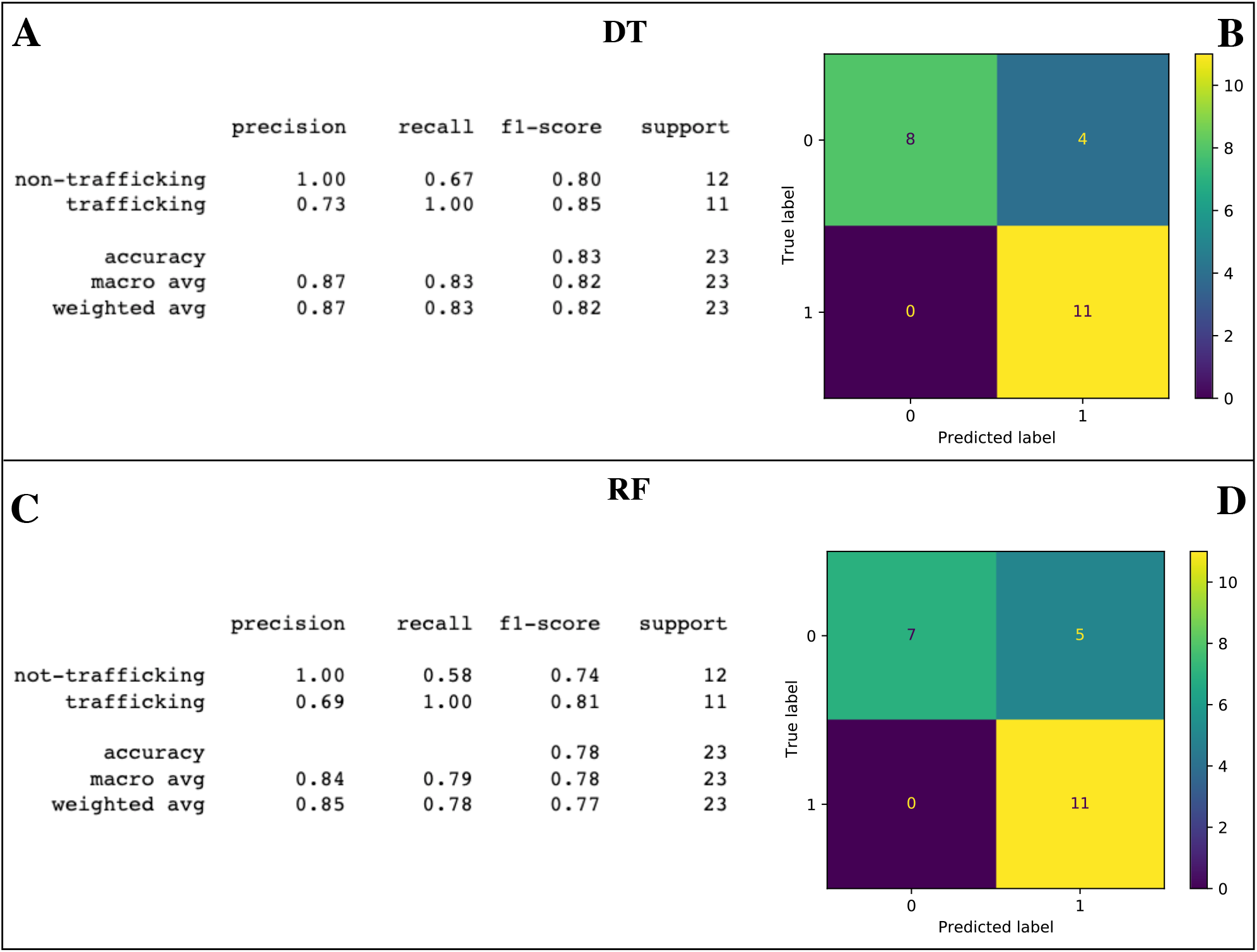
Performance of DT and RF on the top five features (Conservation, FoldX scores, #waters, #H-Bonds, and turns).

**Figure S21:**
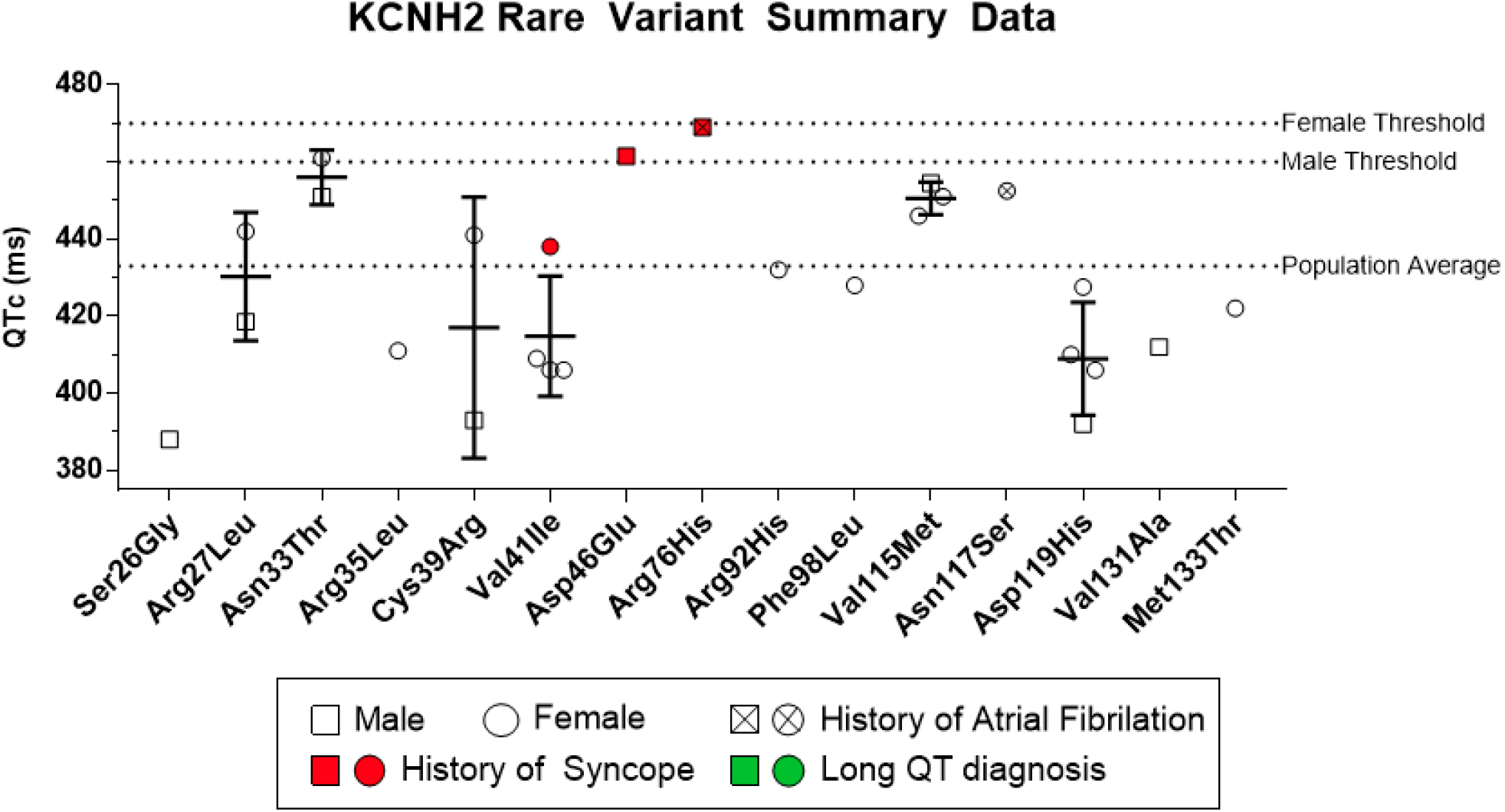
QT intervals measured in patients harboring mutations at PASD sites spanning Ser26-Met133. No patients were diagnosed with long QT and the majority of patients presented sub-threshold QT intervals. Two patients (D46E and R76H) exhibit QT intervals of 460 ms and 470 ms that are commensurate with clinical thresholds for males and females, respectively. Each of these patients has a history of syncope or atrial fibrilation that complicates a clear long QT diagnosis.

**Table S1:**
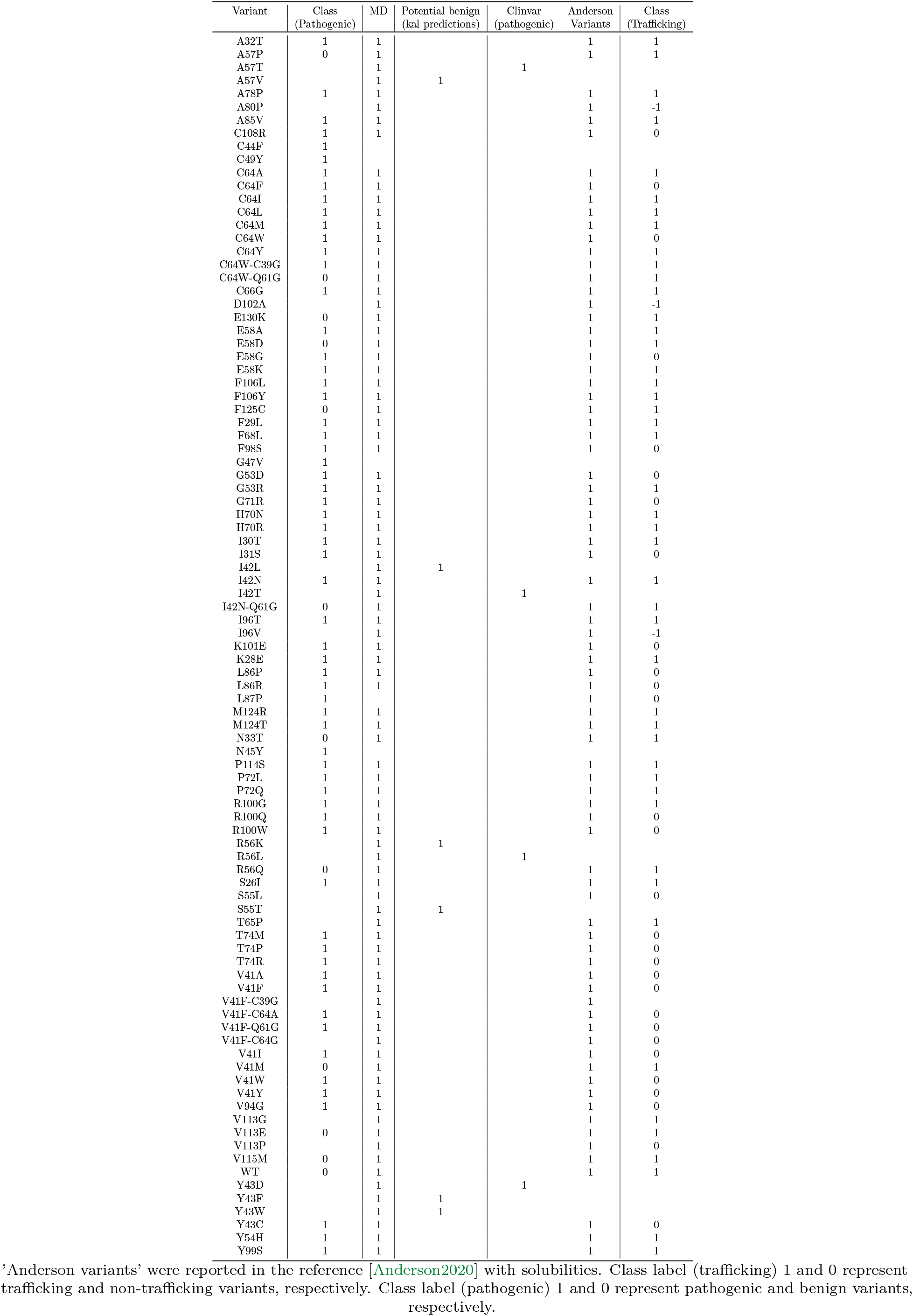
All PAS variants included in this study

